# Functional cross-talk between allosteric effects of activating and inhibiting ligands underlies PKM2 regulation

**DOI:** 10.1101/378133

**Authors:** Jamie A. Macpherson, Alina Theisen, Laura Masino, Louise Fets, Paul C. Driscoll, Vesela Encheva, Ambrosius P. Snijders, Stephen R. Martin, Jens Kleinjung, Perdita E. Barran, Franca Fraternali, Dimitrios Anastasiou

## Abstract

Allosteric regulation is central to the role of the glycolytic enzyme pyruvate kinase M2 (PKM2) in cellular metabolism. Multiple activating and inhibitory allosteric ligands regulate PKM2 activity by controlling the equilibrium between high activity tetramers and low activity dimers and monomers. However, it remains elusive how allosteric inputs upon simultaneous binding of different ligands are integrated to regulate PKM2 activity. Here, we show that, in the presence of the allosteric inhibitor L-phenylalanine (Phe), the activator fructose 1,6-bisphosphate (FBP) can induce PKM2 tetramerisation, but fails to maximally increase enzymatic activity. Guided by a new computational framework we developed to identify residues that mediate FBP-induced allostery, we generated two PKM2 mutants, A327S and C358A, in which activation by FBP remains intact but cannot be attenuated by Phe. Our findings demonstrate a role for residues involved in FBP-induced allostery in enabling the integration of allosteric input from Phe and reveal a mechanism that underlies the co-ordinate regulation of PKM2 activity by multiple allosteric ligands.

---

Allostery refers to the regulation of protein function resulting from the binding of an effector to a site distal to the protein’s functional centre (the active site in the case of enzymes) and is a crucial mechanism for the control of multiple physiological processes^1,2^. The functional response of enzymes to allosteric ligands is thought to occur on the ns - ms time scale, preceding other important regulatory mechanisms such as changes in gene expression or signalling-induced post-translational modifications (PTMs)^3^, thereby enabling fast cellular responses to various stimuli. Despite the demonstrated importance of protein allostery, the investigation of underlying mechanisms remains a challenge for conventional structural approaches and necessitates multi-disciplinary strategies. Latent allosteric pockets can emerge as a consequence of protein flexibility^4,5^ making their identification elusive. Even for known allosteric pockets, an understanding of the molecular mechanisms that underpin the propagation of free energy between an identified allosteric site and the active site can be complicated because of the involvement of protein structural motions on a variety of time scales^6^. Furthermore, many proteins contain several allosteric pockets that facilitate the simultaneous binding of several ligands^7–9^. Concurrent binding of multiple allosteric ligands can modulate the functional response of a protein through the action of multiple allosteric pathways^10^, however, it remains unclear whether such allosteric pathways operate independently, or integrate, either synergistically or antagonistically, to control protein function.

Altered allosteric regulation has a prominent role in the control of tumour metabolism, as well as a number of other pathological processes^2,8,11^. Changes in glycolysis observed in some tumours have been linked to aberrant allosteric regulation of glycolytic enzymes including phosphofructokinase, triose phosphate isomerase and pyruvate kinase. Pyruvate kinases (PKs) catalyse the transfer of a phosphate from phosphoenolpyruvate (PEP) to adenosine diphosphate (ADP) and produce pyruvate and adenosine triphosphate (ATP). There are four mammalian isoforms of pyruvate kinase: PKM1, PKM2, PKL and PKR. PKM2 is highly expressed in tumour cells and in many proliferative tissues, and has critical roles in cancer metabolism that remain under intense investigation^12–15^, as well as in controlling systemic metabolic homeostasis and inflammation^16–18^.

In contrast to the highly homologous alternatively spliced variant PKM1, which is thought to be constitutively active, PKM2 activity in cancer cells is maintained at a low level by the action of various allosteric ligands or post-translational modifications (PTMs)^19^. Low PKM2 activity promotes pro-tumorigenic functions, including divergence of glucose carbons into biosynthetic and redox-regulating pathways that support proliferation and defence against oxidative stress^13,14,20,21^. Small-molecule activators, which render PKM2 constitutively active by overcoming endogenous inhibitory cues, attenuate tumour growth, suggesting that regulation of PKM2 activity, rather than increased PKM2 expression *per se*, is important^13,20,22,23^. Therefore, PKM2 has emerged as a prototypical metabolic enzyme target for allosteric modulators and this has contributed to the renewed impetus to develop allosteric drugs for metabolic enzymes, which are expected to specifically interfere with cancer cell metabolism while sparing normal tissues^11^.

The structure of the PKM2 protomer comprises N-terminal, A, B and C domains. Various ligands that regulate PKM2 activity bind to sites distal from the catalytic core, nestled between the A and B domains. The upstream glycolytic intermediate fructose-1,6-bisphosphate (FBP) binds to a pocket in the C-domain^24^ and increases the enzymatic activity of PKM2, thereby establishing a feed-forward loop that prepares lower glycolysis for increased levels of incoming glucose carbons. Activation of PKM2 by FBP is associated with a decreased K_M_ for the substrate PEP, while the k_cat_ remains unchanged^25–28^, although some reports also find an elevated k_cat_^29,30^ and the reason for this discrepancy remains unknown. Additionally, several amino acids regulate PKM2 activity by binding to a pocket in the TIM-barrel core^25,31,32^. L-serine (Ser) and L-histidine (His) increase, whereas L-phenylalanine (Phe), L-alanine (Ala), L-tryptophan (Trp), L-valine (Val) and L-proline (Pro) decrease the apparent affinity for PEP. Similar to FBP, the reported effects on k_cat_ vary^25,29,31,33^. Furthermore, succinyl-5-aminoimidazole-4-carboxamide-1-ribose 5’-phosphate (SAICAR)^34^ and the triiodothyronine (T3) hormone^29^ bind to unidentified PKM2 pockets to increase and decrease, respectively, the affinity for PEP.

Many of these allosteric effectors have been shown to regulate PKM2 activity by changing the equilibrium of PKM2 between one of three states – a low activity monomer, a low activity tetramer (T-state), and a high activity tetramer (R-state) – with some studies reporting the existence of low activity dimers^24,29,35–39^. FBP promotes, whereas T_3_ prevents, tetramerisation^24,29,40^. The mode of PKM2 regulation by amino acids is unclear. Some reports suggest that Ala promotes the formation of inactive PKM2 dimers^38,41^, whereas others show that Phe, Ala and Trp stabilise the T-state tetramer^29,31^. In addition to allosteric effectors, PTMs can also influence the oligomerisation of PKM2 protomers, although the effects of PTMs on the enzyme mechanism remain elusive. Nevertheless, PKM2 activity in cells is often inferred from the oligomeric state PKM2 is found to adopt^13,18,20,22,42–46^. Collectively, current evidence suggests that a link between enzyme activity and oligomerisation state exists and, while not well understood, it may play a role in PKM2 regulation.

*In vitro*, PKM2 can bind concurrently to multiple allosteric effectors that might either reciprocally influence each other’s action or exert independent effects on enzymatic activity. A PKM2 mutant that cannot bind FBP can still be activated by Ser and, conversely, a mutant that abolishes Ser binding can be activated by FBP^25^, suggesting that amino acids could work independently from FBP to regulate PKM2. However, inhibitory amino acids that bind to the same pocket as Ser fail to inhibit the enzyme in the presence of FBP indicating a dominant influence of the latter^31,47^. Similarly, FBP can overcome PKM2 inhibition by T_3_^40^. FBP has also been shown to attenuate, but not completely prevent, inhibition of PKM2 by Ala^48^. These observations raise the question of how inputs from multiple cues are integrated to regulate PKM2 activity. This issue is particularly pertinent in cells, where multiple allosteric ligands coexist at varying concentrations and bind PKM2 with a range of affinities.

Here we provide evidence that Phe can bind simultaneously with FBP to PKM2 in cells, and show that Phe attenuates activation of purified PKM2 by FBP in the absence of large tertiary or quaternary structure changes. Using a novel computational framework to predict residues implicated in allosteric signal transmission from molecular dynamics (MD) simulations, we identify a network of PKM2 residues that mediates allosteric activation by FBP. Intriguingly, mutagenesis of some of these residues interferes with the ability of Phe to hinder PKM2 regulation by FBP but does not perturb FBP–induced activation itself. These residues, therefore, integrate inputs from distinct allosteric effectors and underlie the functional synergism between ligands that bind to distinct pockets in regulating PKM2 activity.

## RESULTS

### Simultaneous binding of multiple ligands to PKM2 is relevant for its regulation in cells

The fraction of PKM2 bound to its allosteric effectors in cells, and hence the ability of these ligands to regulate PKM2 activity, is determined by their respective binding affinities and intracellular concentrations. To explore whether FBP and amino acids are likely to bind simultaneously and whether this is relevant for the regulation of PKM2, we aimed to assess the fractional saturation of PKM2 bound to FBP, Phe and Ser, in proliferating cancer cells.

To this end, we first measured the affinity of PKM2 for FBP, Phe and Ser *in vitro*, using fluorescence emission spectroscopy and microscale thermophoresis. Similar to previous reports^35,46^, we found that purified recombinant PKM2 remained bound to FBP, to varying degrees (**Supplementary Fig. 1a-d**), despite extensive dialysis, suggesting a high binding affinity. Consistent with this prediction, the K_D_^FBP^ for PKM2 was (21.4 ± 9.0) nM, after accounting for the amount of co-purified FBP (**Supplementary Fig. 1e** and **Table 1**). The K_D_^Ser^ and K_D_^Phe^ were (507.5 ± 218.2) μM and (191.0 ± 86.3) μM, respectively (**Supplementary Fig. 1f** and **Table 1**).

**Table 1.** Apparent steady-state binding constants of PKM2 for FBP, Phe and Ser.

| Titrated ligand | Competing ligand | $K_D^{\text{apparent}}$ |
| --- | --- | --- |
| FBP | – | (21.4 ± 9.0) nM |
|  | Ser | (12.5 ± 7.7) nM |
|  | Phe | (8.1 ± 5.6) nM |
| Phe | – | (191.0 ± 86.3) μM |
|  | FBP | (132.0 ± 11.7) μM |
| Ser | – | (507.5 ± 218.2) μM |
|  | FBP | (462.0 ± 97.1) μM |

We next calculated the predicted fractional saturation range of PKM2 with FBP, Ser and Phe in cells. To this end, we determined the intracellular concentration of PKM2, using targeted proteomics, and the range of intracellular concentrations of FBP, Ser and Phe (denoted as [X]_ic_, where X is the respective metabolite), using metabolomics, in three human cancer cell lines (see Methods). In the presence of high (11 mM) extracellular glucose, [FBP]_ic_ varied between 240-360 μM across the three cell lines (Gluc^+^, **Fig. 1a** and **Table 2**), a concentration range between 12000– and 18000–fold in excess of its binding affinity to PKM2. The calculated fractional saturation of PKM2 with FBP was 0.99 (**Fig. 1b**) and remained unchanged even in cells cultured in the absence of glucose (Gluc^−^), when [FBP]_ic_ decreased to as low as 20 μM. In the presence of near-physiological extracellular concentrations of Phe or Ser (Phe^100^ or Ser^100^), [Phe]_ic_ and [Ser]_ic_ across the three lines ranged between 220 – 580 μM for Phe and 2000 – 6000 μM for Ser (**Fig. 1c** and **Table 2**), close to their respective binding affinities for PKM2. The predicted fractional saturation range was 0.53 – 0.75 and 0.81 – 0.92 for Phe and Ser, respectively (**Fig. 1d, e**). The range of predicted fractional saturations in complete absence of amino acids (-aa) or 5x physiological concentrations was 0.05 (-aa) to 0.90 (Phe^500^) for Phe (**Fig. 1d**) and 0.18 (-aa) to 0.98 (Ser^500^) for Ser (**Fig. 1e**). Together, these calculations predict that, in a range of relevant extracellular nutrient concentrations, FBP binding to PKM2 is near-saturating, whereas that of amino acids is not.

**Figure 1:**
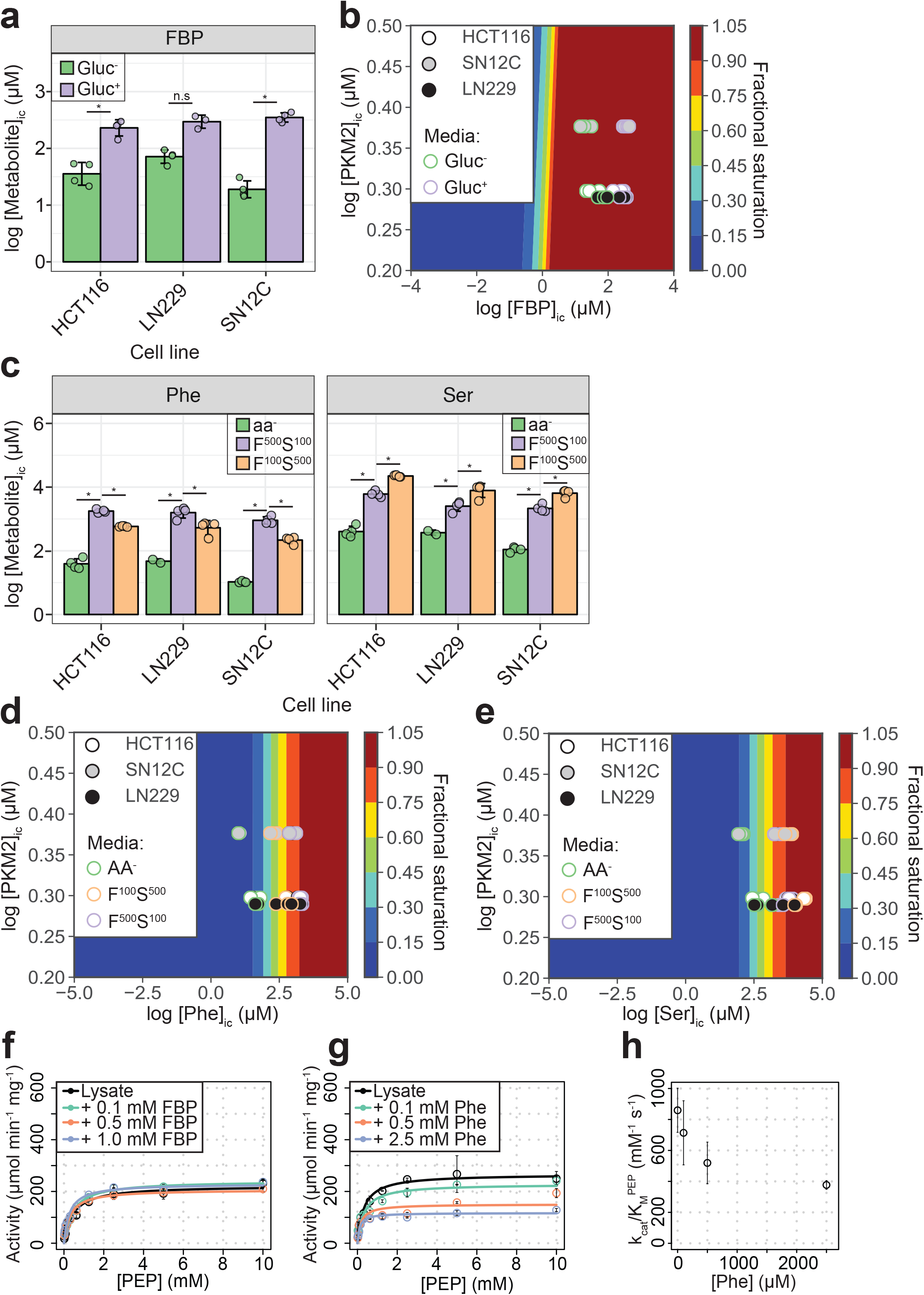
PKM2 allosteric effector concentrations in cells predict saturating binding of FBP and sub-saturating binding of Phe and Ser. **(a)** Intracellular concentrations of FBP ([FBP]_ic_) measured using liquid-chromatography mass spectrometry (LC-MS), from HCT116 (colorectal carcinoma), LN229 (glioblastoma) and SN12C (renal cell carcinoma) cells cultured in RPMI media containing 11 mM glucose (Gluc^+^), or 0 mM (Gluc^−^) for 1 h. Statistical significance was assessed using a Wilcoxon rank-sum test. Asterisk (*) marks significant changes (p-value < 0.05). **(b)** Phase diagram for intracellular FBP binding to PKM2 computed for a range of [FBP] and [PKM2] values using the binding constant obtained as shown in **Supplementary Fig. 1e.** Colour scale represents fractional saturation of PKM2 with ligand. A fractional saturation of 0 indicates no FBP bound to PKM2 and a fractional saturation equal to 1 indicates that each FBP binding site in the cellular pool of PKM2 would be occupied. Experimental fractional saturation values were estimated from [FBP]_ic_ obtained from **(a)** and [PKM2]_ic_ was determined using targeted proteomics (see Methods and **Supplementary Table**). The predicted fractional saturation for each of the three cell lines (four technical replicates) is shown as shaded open circles in the phase diagram. **(c)** Intracellular concentrations of Phe ([Phe]_ic_) and Ser ([Ser]_ic_) measured as in **(a)**, in HCT116, LN229 and SN12C cells cultured in Hank’s Balanced Salt Solution (HBSS) without amino acids (aa^−^), HBSS containing 100 μM Phe and 500 μM Ser (F^100^ S^500^), or HBSS containing 500 μM Phe and 100 μM Ser (F^500^ S^100^). The low concentrations of Phe and Ser are similar to human serum concentrations^74^. [Phe]_ic_ and [Ser]_ic_ were not affected by extracellular glucose concentration, neither did extracellular Phe and Ser concentrations influence [FBP]ic (**Supplementary Fig. 2a, b**). Statistical significance was assessed as in **(a). (d)** Phase diagram for Phe computed as in **(b)** using [Phe]_ic_ from **(c). (e)** Phase diagram for Ser computed as in **(b)** using [Ser]ic from **(c). (f)** PKM2 activity in lysates of HCT116 cells cultured in RPMI (Gluc^+^). Measurements were repeated following the addition of either 0.1, 0.5 or 1.0 mM of exogenous FBP. Initial velocity curves were fitted using Michaelis-Menten kinetics and the absolute concentration of PKM2 in the lysates was estimated using quantitative Western blotting (**Supplementary Fig. 2d**), to calculate PKM2 specific activity. **(g)** PKM2 activity in HCT116 cell lysates as in **(f)**, but with addition of exogenous Phe. **(h)** Plot of k_cat_/K_M_ versus [Phe] from **(g)** revealing a dose-dependent inhibitory effect of Phe on the activity of PKM2 in HCT116 lysates.

**Table 2.** Intracellular concentrations of FBP, Phe and Ser in cancer cell lines.

| Metabolite | Media condition | Cell line | [Metabolite]<sub>ic</sub><br>(μM) | Fractional saturation |
| --- | --- | --- | --- | --- |
| FBP | Gluc<sup>–</sup><br>(RPMI) | HCT116 | 38.4 ± 16.6 | 0.99 ± 0.00 |
|  |  | LN229 | 73.4 ± 18.4 | 0.99 ± 0.00 |
|  |  | SN12C | 19.9 ± 7.5 | 0.98 ± 0.00 |
|  | Gluc<sup>+</sup><br>(RPMI) | HCT116 | 238.4 ± 67.4 | 0.99 ± 0.00 |
|  |  | LN229 | 302.0 ± 75.3 | 0.99 ± 0.00 |
|  |  | SN12C | 355.1 ± 66.2 | 0.99 ± 0.00 |
|  | Gluc<sup>+</sup> aa<sup>–</sup><br>(HBSS) | HCT116 | 337.2 ± 185.5 | 0.99 ± 0.00 |
|  |  | LN229 | 257.0 ± 33.9 | 0.99 ± 0.00 |
|  |  | SN12C | 236.8 ± 41.1 | 0.99 ± 0.00 |
|  | Gluc<sup>+</sup> Phe<sup>500</sup> Ser<sup>100</sup><br>(HBSS) | HCT116 | 384.5 ± 41.8 | 0.99 ± 0.00 |
|  |  | LN229 | 277.3 ± 83.4 | 0.99 ± 0.00 |
|  |  | SN12C | 300.3 ± 89.1 | 0.99 ± 0.00 |
|  | Gluc<sup>+</sup> Phe<sup>100</sup> Ser<sup>500</sup><br>(HBSS) | HCT116 | 400.9 ± 82.2 | 0.99 ± 0.00 |
|  |  | LN229 | 372.2 ± 106.0 | 0.99 ± 0.00 |
|  |  | SN12C | 286.4 ± 73.6 | 0.99 ± 0.00 |

| Metabolite | Media condition | Cell line | [Metabolite]<sub>ic</sub><br>(μM) | Fractional<br>saturation |
| --- | --- | --- | --- | --- |
| Phe | Gluc<sup>-</sup><br>(RPMI) | HCT116 | 256.3 ± 65.8 | 0.56 ± 0.06 |
| | | LN229 | 231.6 ± 22.6 | 0.55 ± 0.03 |
| | | SN12C | 100.4 ± 24.6 | 0.34 ± 0.06 |
| | Gluc<sup>+</sup><br>(RPMI) | HCT116 | 258.2 ± 79.6 | 0.56 ± 0.08 |
| | | LN229 | 318.7 ± 115.8 | 0.61 ± 0.09 |
| | | SN12C | 197.5 ± 26.6 | 0.51 ± 0.04 |
| | Gluc<sup>+</sup> aa<sup>-</sup><br>(HBSS) | HCT116 | 41.1 ± 15.8 | 0.17 ± 0.05 |
| | | LN229 | 31.7 ± 28.0 | 0.13 ± 0.12 |
| | | SN12C | 10.6 ± 0.6 | 0.05 ± 0.02 |
| | Gluc<sup>+</sup> Phe<sup>500</sup> Ser<sup>100</sup><br>(HBSS) | HCT116 | 1776.2 ± 225.1 | 0.90 ± 0.01 |
| | | LN229 | 1666.5 ± 543.0 | 0.89 ± 0.04 |
| | | SN12C | 926.1 ± 252.7 | 0.82 ± 0.04 |
| | Gluc<sup>+</sup> Phe<sup>100</sup> Ser<sup>500</sup><br>(HBSS) | HCT116 | 582.4 ± 23.8 | 0.75 ± 0.01 |
| | | LN229 | 575.7 ± 224.4 | 0.73 ± 0.11 |
| | | SN12C | 221.6 ± 49.1 | 0.53 ± 0.06 |
| Ser | Gluc<sup>-</sup><br>(RPMI) | HCT116 | 3580.9 ± 1016.9 | 0.87 ± 0.03 |
| | | LN229 | 1755.0 ± 159.5 | 0.77 ± 0.02 |
| | | SN12C | 854.3 ± 240.7 | 0.62 ± 0.07 |
| | Gluc<sup>+</sup><br>(RPMI) | HCT116 | 3943.5 ± 1363.1 | 0.88 ± 0.05 |
| | | LN229 | 2226.7 ± 757.7 | 0.80 ± 0.06 |
| | | SN12C | 1766.0 ± 142.9 | 0.78 ± 0.01 |
| | Gluc<sup>+</sup> aa<sup>-</sup><br>(HBSS) | HCT116 | 426.9 ± 179.2 | 0.44 ± 0.09 |
| | | LN229 | 249.6 ± 221.8 | 0.28 ± 0.25 |
| | | SN12C | 111.0 ± 18.4 | 0.18 ± 0.02 |
| | Gluc<sup>+</sup> Phe<sup>500</sup> Ser<sup>100</sup><br>(HBSS) | HCT116 | 6157.3 ± 1334.4 | 0.92 ± 0.01 |
| | | LN229 | 2641.1 ± 825.1 | 0.83 ± 0.06 |
| | | SN12C | 2197.7 ± 605.8 | 0.81 ± 0.04 |
| | Gluc<sup>+</sup> Phe<sup>100</sup> Ser<sup>500</sup><br>(HBSS) | HCT116 | 22360.4 ± 1554.2 | 0.98 ± 0.00 |
| | | LN229 | 8464.7 ± 3214.5 | 0.93 ± 0.04 |
| | | SN12C | 6603.9 ± 1500.4 | 0.93 ± 0.02 |

Consistent with the prediction of near-saturating FBP occupancy, addition of exogenous FBP to lysates of HCT116 cells cultured in both Gluc^+^ and Gluc^−^ media caused little increase in PKM2 activity (**Fig. 1f** and **Supplementary Fig. 2c,d**). Addition of exogenous Phe (at physiological intracellular concentrations) to HCT116 lysates resulted in inhibition of PKM2 activity (**Fig. 1g**) and a dose-dependent decrease in k_cat_/K_M_ (**Fig. 1h**). Addition of exogenous Ser to HCT116 lysates reversed the inhibition of PKM2 activity by Phe, consistent with competitive binding between Ser and Phe for the same PKM2 pocket^31^ (**Supplementary Fig. 2d,e**).

Together, these observations support the concept that, during steady-state cell proliferation under typical culture conditions, a significant fraction of PKM2 is bound to FBP. Under these conditions, amino acids can reversibly bind to PKM2, and could thereby regulate PKM2 activity, in the background of pre-bound FBP.

### Phe inhibits FBP-bound PKM2 without causing PKM2 tetramer dissociation

To investigate how allosteric ligands, alone and in combination, regulate PKM2, we first measured their effects on PKM2 enzyme activity and then on its oligomerisation. We found that FBP decreased the K_M_^PEP^ to (0.23 ± 0.04) mM, compared to the absence of any added ligands (PKM2^apo^) [K_M_^PEP^ = (1.22 ± 0.02) mM] (**Fig. 2a** and **Table 3**). Similarly, addition of Ser resulted in a decrease of the K_M_^PEP^ to (0.22 ± 0.04) mM, whereas Phe increased the K_M_^PEP^ to (7.08 ± 1.58) mM. None of the three ligands changed the k_cat_. These results are consistent with previous reports^24,25,49^ that FBP, Phe and Ser change the K_M_^PEP^ but not the k_cat_ and therefore act as K-type modulators^50^ of PKM2.

**Figure 2:**
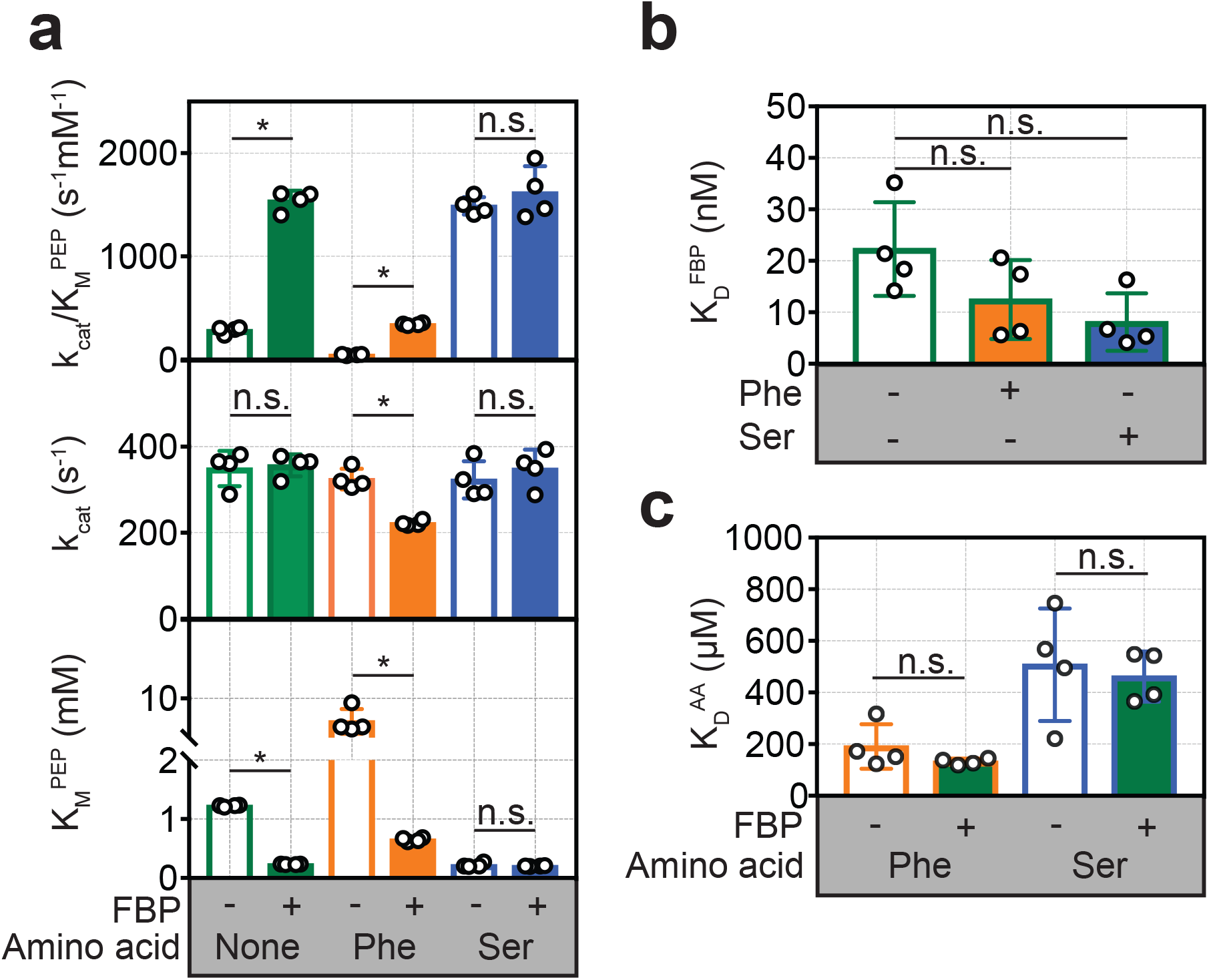
FBP influences the inhibition of PKM2 activity by Phe. **(a)** Steady-state kinetic parameters of purified recombinant human PKM2, in the absence of added ligands; in the presence of 2 μM FBP, 400 μM Phe and 200 mM Ser alone; and after addition of either 400 μM Phe or 200 mM Ser to PKM2 pre-incubated with 2 μM FBP. Initial velocity curves were fit to Michaelis-Menten kinetic models. Each titration was repeated four times. Statistical significance was assessed using a Wilcoxon rank-sum test. Asterisk (*) marks significant changes (p-value < 0.05). **(b)** Binding constant of FBP to PKM2 obtained from fluorescence emission spectroscopy measurements in the absence and in the presence of either 400 μM Phe or 200 mM Ser. **(c)** Binding constants for Phe and Ser to PKM2, in the absence or presence of 2 μM FBP, obtained from microscale thermophoresis (MST) measurements. MST was used, rather than intrinsic fluorescence spectroscopy as for FBP, due to the absence of tryptophan residues proximal to the amino acid binding pocket on PKM2. Significance was assessed as in **(a)**.

**Table 3.** Steady-state Michaelis-Menten kinetic parameters for PKM2^apo^, and following the addition of FBP, Phe and Ser.

| **PKM2<sup>ligand</sup>** | **K<sub>M</sub><sup>PEP</sup> (mM)** | **k<sub>cat</sub> (s<sup>-1</sup>)** | **k<sub>cat</sub>/K<sub>M</sub><sup>PEP</sup> (s<sup>-1</sup> mM<sup>-1</sup>)** |
| --- | --- | --- | --- |
| PKM2<sup>apo</sup> | 1.22 ± 0.02 | 349.3 ± 40.9 | 285.6 ± 34.1 |
| PKM2<sup>Phe</sup> | 7.08 ± 1.58 | 324.7 ± 23.9 | 46.8 ± 6.0 |
| PKM2<sup>Ser</sup> | 0.22 ± 0.04 | 323.1 ± 43.2 | 1489.8 ± 84.7 |
| PKM2<sup>FBP</sup> | 0.23 ± 0.04 | 356.7 ± 25.7 | 1540.4 ± 96.9 |
| PKM2<sup>FBP + Phe</sup> | 0.65 ± 0.03 | 222.3 ± 6.3 | 342.1 ± 11.1 |
| PKM2<sup>FBP + Ser</sup> | 0.20 ± 0.04 | 348.7 ± 44.3 | 1620.0 ± 253.6 |

In the presence of FBP, addition of Phe resulted in a decrease in the k_cat_ of PKM2 from 349.3 s^−1^ to 222.3 s^−1^ (p = 0.0075), and a simultaneous increase in the K_M_^PEP^ [(0.65 ± 0.03) mM] compared to PKM2^FBP^ (**Fig. 2a** and **Supplementary Fig. 3a-h**). Further analysis of the kinetic data showed that in the absence of FBP, Phe acted as a hyperbolic-specific^51^ inhibitor (**Supplementary Fig. 3i**), whereas, with FBP, Phe inhibition of PKM2 changed to a hyperbolic-mixed^51^ mechanism (**Supplementary Fig. 3j**). In contrast, Ser caused no changes to either the K_M_^PEP^ or k_cat_ in the presence of FBP (**Fig. 2a**). Given that Phe and Ser bind to the same pocket^31^, we focused on Phe as an amino acid modulator of PKM2 for further investigations into the functional interaction between the amino acid and FBP binding pockets.

To explore the possibility that Phe modifies FBP binding and *vice versa*, we measured the binding affinities of each ligand alone, or in the presence of the other. Phe caused a small but not statistically significant (p = 0.150) increase in the binding affinity of FBP (K_D_^FBP^) (**Fig. 2b**), and, conversely, saturating amounts of FBP did not change the measured K_D_^Phe^ (**Fig. 2c**). These measurements suggested that decreased binding of FBP cannot account for the attenuation of FBP–induced activation of PKM2 upon addition of Phe.

We next used nano-electrospray ionisation (nESI) and ion mobility (IM) mass spectrometry (MS)^52,53^ to investigate whether Phe impedes FBP–induced activation of PKM2 by perturbing the oligomeric state of the protein. The mass spectrum of PKM2^apo^ showed a mixture of monomers, dimers and tetramers at an approximate ratio of 1:7:10 (**Fig. 3a,b**), with dimers exclusively forming the A-A’ dimer assembly, as evidenced from experimental (IM) and theoretical measurements (**Supplementary Fig. 4a,b**). Mass-deconvolution of the spectrum of PKM2^apo^ indicated that the signal of the tetramer consisted of five distinct species, which were assigned to PKM2^apo^; and PKM2 bound to 1, 2, 3 or 4 molecules of FBP (**Fig. 3c**), consistent with our earlier finding that FBP co-purifies with PKM2 (**Supplementary Fig. 1d**). Addition of exogenous FBP resulted in a decrease in the intensity of monomer and dimer peaks, and a dose-dependent increase in the intensity of the tetramer signal (**Supplementary Fig. 4c,d**). At saturating concentrations of FBP, the mass spectrum of PKM2 consisted of a single species corresponding to tetrameric PKM2 bound to four molecules of FBP (**Fig. 3c**). Furthermore, surface-induced dissociation (SID) experiments revealed that PKM2^FBP^ tetramers were more stable than PKM2^apo^ tetramers with respect to dissociation into dimers and monomers (**Supplementary Fig. 5**). Together, these data indicated that addition of FBP promotes the formation of PKM2 tetramers, as seen previously using other methods^37,40^, and confer an enhanced tetramer stability.

**Figure 3:**
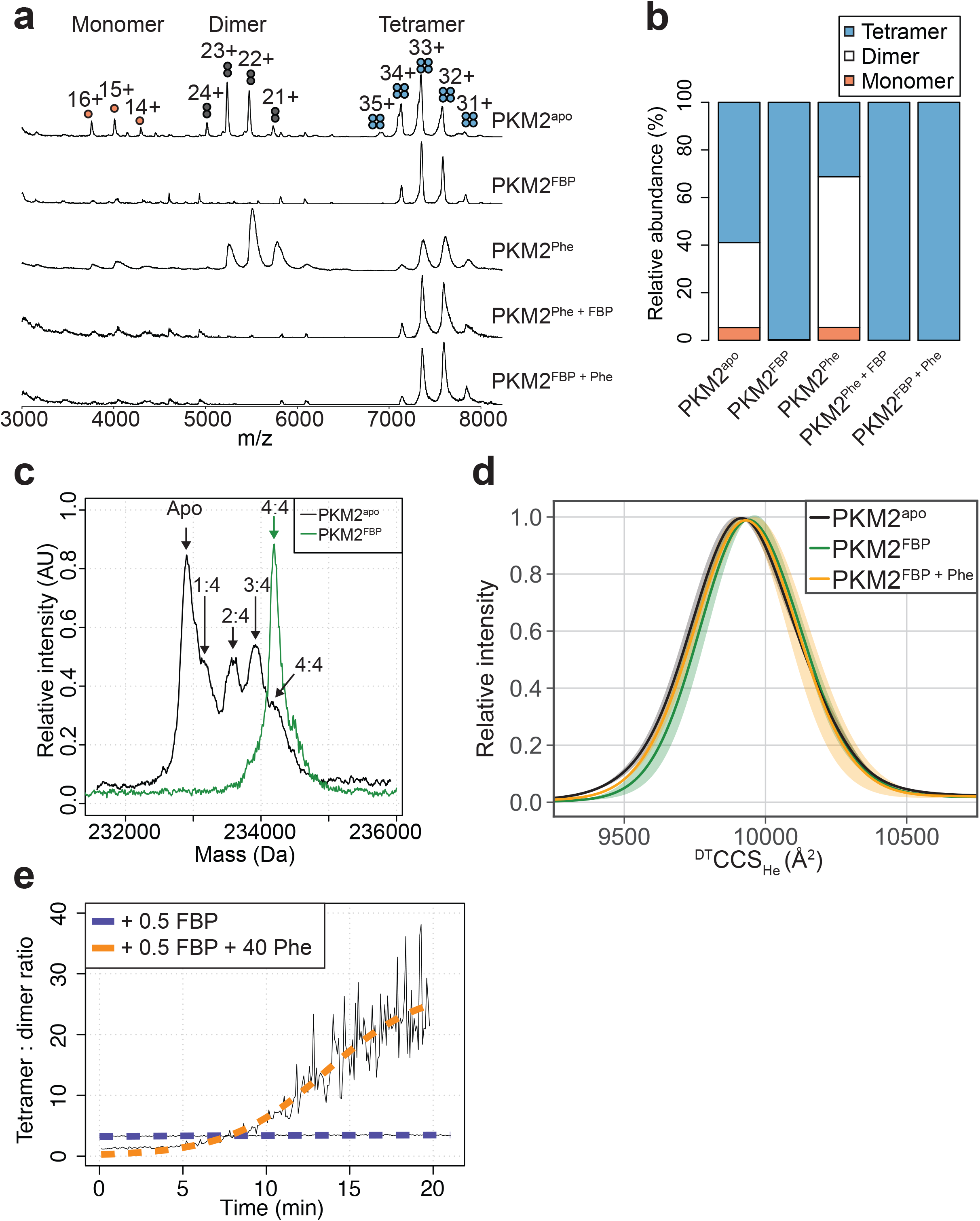
FBP modulates the effects of Phe on PKM2 oligomerisation. **(a)** Native mass spectra of 10 μM PKM2 in 200 mM ammonium acetate at pH 6.8, in the absence of allosteric ligands (PKM2^apo^), or in the presence of: 10 μM FBP (PKM2^FBP^), 300 μM Phe (PKM2^Phe^), 300 μM Phe followed by addition of 10 μM FBP (PKM2^Phe + FBP^) or 10 μM FBP followed by addition of 300 μM Phe (PKM2^FBP + Phe^). **(b)** Relative abundance of PKM2 monomers, dimers and tetramers obtained from the spectra shown in (a) by computing the area of the peaks corresponding to each of the three oligomeric states. Relative peak areas were calculated as a percentage of the total area given by all charge-state species in a single mass spectrum. **(c)** Deconvolved mass spectra of PKM2 tetrameric species in the absence of any added ligands (PKM2^apo^) or presence of FBP (PKM2^FBP^). pKM2^apo^ has five distinct mass peaks, separated by 340 Da (equivalent to the weight of FBP), corresponding to tetrameric PKM2^apo^, and tetrameric PKM2 bound to 1, 2, 3 and 4 molecules of FBP, respectively. The spectrum of PKM2^FBP^ contains a single peak, corresponding to tetrameric PKM2 bound to four molecules of FBP. **(d)** ^DT^CCS_He_ distribution of PKM2^apo^, PKM2^FBP^ and PKM2^FBP+Phe^ tetramers calculated from analyses of arrival time distribution measurements of PKM2 tetramer peaks (see Methods). **(e)** Change in the oligomeric state of PKM2 over time. Oligomerisation is reported as the ratio of the tetramer 33+ charge-state peak relative to the dimer 22+ charge-state peak, obtained from mass spectra of 10 μM PKM2 following addition of either 5 μM FBP, or 5 μM FBP and 400 μM Phe over the course of 20 min. The kinetics of tetramerisation were estimated from a two-state sigmoidal model (orange and blue dashed lines, see Methods).

Addition of Phe to PKM2^apo^ caused a decrease in the relative abundance of the tetramers and an increased relative abundance of the dimer species (**Fig. 3a,b**). In contrast, Phe did not perturb the tetrameric state of PKM2^FBP^ (**Fig. 3a,b**), while FBP binding to PKM2^Phe^ resulted in tetramerisation of the protein, indicating that the dominant effect of FBP on PKM2 tetramerisation was unaffected by the order of ligand addition (**Fig. 3a,b**). PKM2 tetramers were found to bind FBP and Phe simultaneously, on the basis of the charge-state shift resulting from sequential addition of either ligand alone or in combination (**Supplementary Fig. 6a,b**). Additionally, differences in the collision cross section (^DT^CCS_He_) distribution, which reflects the conformational heterogeneity of proteins, suggested that FBP caused subtle conformational changes in the tetramer that were partially reversed upon subsequent addition of Phe (**Fig. 3d**). Moreover, in the presence of Phe, half-stoichiometric amounts of FBP were sufficient to induce tetramerisation with slow kinetics [*k*_tet_ = (812.5 s ± 284.6 s^−1^)], whereas equivalent half-stoichiometric amounts of FBP in the absence of Phe were unable to fully convert PKM2 monomers and dimers into tetramers (**Fig. 3e**). The propensity for Phe to enhance FBP-induced tetramerisation indicates a functional synergism between the two allosteric ligands, that favours tetramer formation despite the opposing effects of these ligands, individually, both on activity and oligomerisation.

Together, these data demonstrate that the inhibitory effect of Phe on the ability of FBP to enhance PKM2 activity is not due to Phe preventing FBP–induced tetramerisation and suggest it is likely due to interference with the allosteric communication between the FBP and the active site.

### Molecular dynamics simulations reveal candidate residues that mediate FBP–induced PKM2 allostery

To gain insight into the mechanism by which Phe interferes with FBP–induced allosteric activation of PKM2, we first sought to identify PKM2 residues that mediate the allosteric communication between the FBP binding site and the catalytic centre. To this end, we performed molecular dynamics (MD) simulations of PKM2^apo^ tetramers and PKM2^FBP^ tetramers (**Table 4**). Subsequent analyses of protein volume and solvent accessibility of the trajectories (**Supplementary Fig. 7a,b**), found no evidence of global protein structural changes induced by FBP. Furthermore, we found no significant difference in the time-dependent configurational entropy of the PKM2^apo^ and PKM2^FBP^ simulated trajectories (**Supplementary Fig. 7c**), negating the likelihood of a purely entropy-driven allosteric mechanism. We therefore reasoned that enthalpic motions likely play a role in the allosteric regulation of PKM2 by FBP.

**Table 4.** Summary of molecular dynamics simulations.

| **Liganded state simulated** | **PDB ID of starting structure** | **Number of protein atoms** | **Number of water atoms** | **Simulation time** | **Number of replicas** |
| --- | --- | --- | --- | --- | --- |
| PKM2<sup>apo</sup> | 3BJT | 20104 | 263384 | 400 ns | 5 |
| PKM2<sup>FBP</sup> | 3BJF | 20122 | 261594 | 420 ns | 5 |

To test this hypothesis, we set out to identify whether FBP elicits correlated motions^54^ in the backbone of PKM2, and whether these concerted motions form the basis of a network of residues that connect the allosteric pocket to the active site. Accurately computing the network of correlated protein motions from MD simulations is complicated by the occurrence of dynamic conformational sub-states that can display unique structural properties^55,56^. We therefore developed a novel computational framework, named *AlloHubMat* (**Allo**steric **Hub** prediction using **Mat**rices that capture allosteric coupling), to predict allosteric hub fragments from the network of dynamic correlated motions, based on explicitly identified conformational sub-states from multiple MD trajectories (**Fig. 4a**). Extraction of correlated motions from multiple substates within a consistent information theoretical framework allowed us to compare the allosteric networks, both between replicas of the same liganded state and between different liganded states of PKM2 (see Methods).

**Figure 4:**
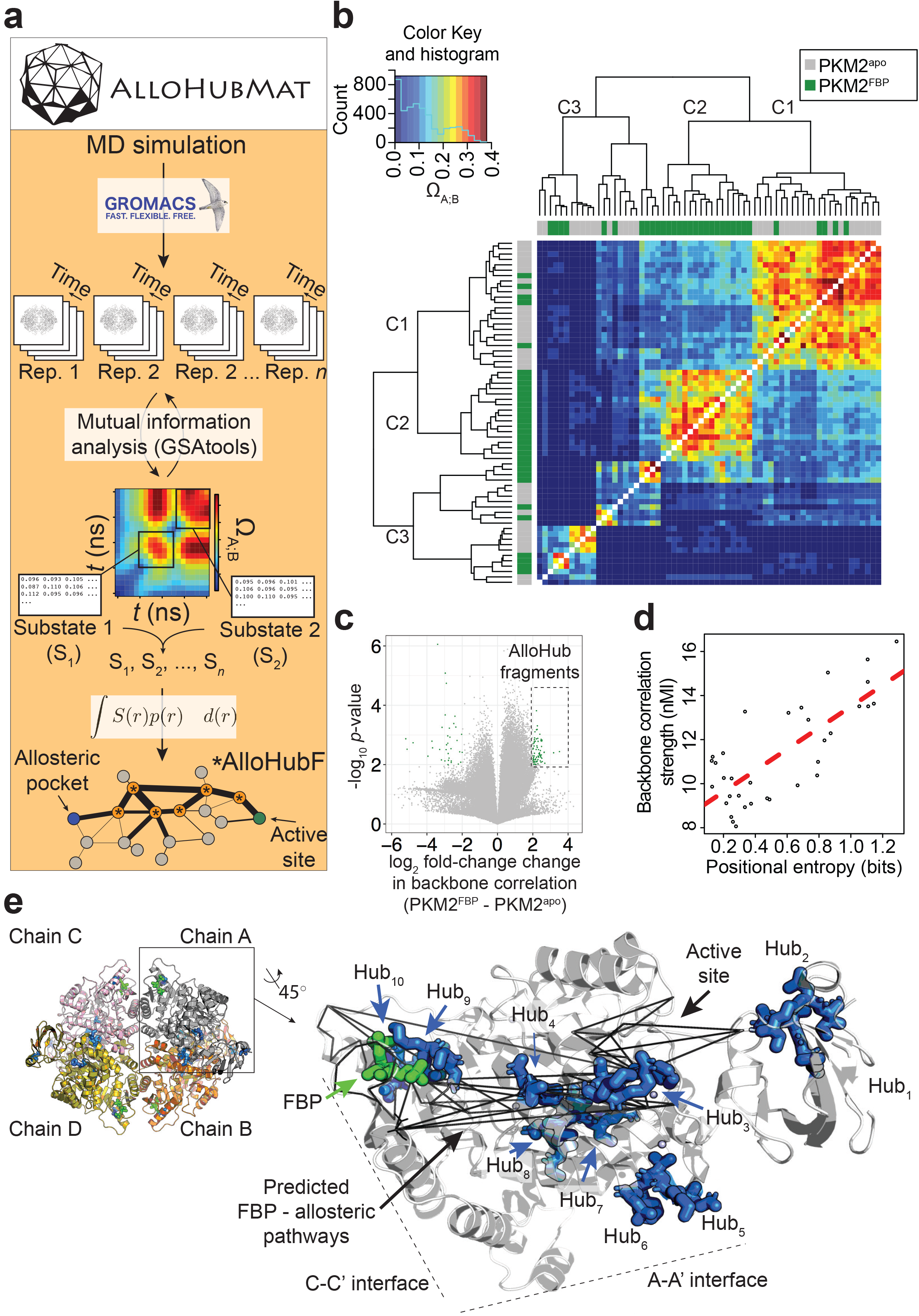
AlloHubMat predicts candidate residues that mediate the allosteric effect of FBP on PKM2, from molecular dynamics (MD) simulations. **(a)** Schematic of the AlloHubMat computational pipeline, developed to identify residues that are involved in the transmission of allostery between an allosteric ligand pocket and the active site. Multiple replicate molecular dynamics (MD) simulations are seeded from the 3D protein structure using the GROMACS molecular dynamics engine. All MD simulations are encoded with the M32K25 structural alphabet^61^, and the protein backbone correlations over the MD trajectory are computed with GSAtools^63^ using information theory mutual information statistics. The backbone correlations are explicitly used to identify and extract configurational sub-states from the MD trajectories. A global allosteric network is then constructed by integrating over the correlation matrices, and their respective probabilities, from which allosteric hub fragments (AlloHubFs) are extracted. Each AlloHubF comprises four consecutive amino acid residues. **(b)** Correlation matrices cluster according to the liganded state of PKM2 in the MD simulations. AlloHubMat, described in **(a)**, was used to identify correlation matrices of the conformational substates from five separate 400 ns MD simulations of PKM2^apo^ (grey) and PKM2^FBP^ (green). In total, we identified eight sub-states for all simulations of PKM2^apo^ and eight for PKM2. For every sub-state, the network of correlations from each of the four protomers is presented individually. To investigate whether the correlated motions for each sub-state could be attributed to the liganded state of PKM2, the correlation matrices were compared with a complete-linkage hierarchical clustering (see Methods). The matrix covariance overlap (Ω_A;B_) was used as a distance metric, represented by the colour scale. A high Ω_A;B_ score indicates high similarity between two correlation matrices, and a low Ω_A;B_ score indicates that the correlation matrices are dissimilar. The clustering analysis revealed three clusters, denoted C1-C3. Cluster C1 was dominated by correlation matrices extracted from PKM2^apo^ simulations, and cluster C2 was exclusively occupied by PKM2^FBP^ correlation matrices. C3 consisted of correlation matrices from PKM2^apo^ and PKM2^FBP^ simulations. **(c)** A volcano plot showing difference in protein backbone correlations – derived from the AlloHubMat analysis – between PKM2^apo^ and PKM2^FBP^. Each point corresponds to a correlation between two distal protein fragments; points with a positive log_2_ fold-change represent correlations that are predicted to increase in strength upon FBP binding. Correlations with a log_2_ fold-change ≥ 2 and a false discovery rate ≤ 0.05 % (determined from a Wilcoxon ranked-sum test) between PKM2^apo^ and PKM2 were designated as AlloHub fragments, highlighted in green. A total of 72 AlloHub fragments were predicted from this analysis. **(d)** The positional entropy of the PKM2 fragment-encoded structure correlates linearly with the correlation strength of the fragment. The total mutual information content was computed by summing over the correlations for each of the top AlloHub fragments. nMI: normalised mutual information. **(e)** Left: PKM2 structure depicting the spatial distribution of the top ten predicted AlloHub fragments. Right: zoom into a single protomeric chain shown in cartoon representation. AlloHub fragments (blue) and FBP (green) are shown as stick models. Black lines indicate minimal distance pathways between the FBP binding pocket and the active site, predicted using Dijkstra’s algorithm (see Methods) with the complete set of correlation values as input.

Using AlloHubMat, we analysed all replicate MD simulations of PKM2^apo^ and PKM2^FBP^ and identified several conformational sub-states. Backbone correlations extracted from the substates separated into two clusters (C1 and C2) that were dominated by sub-states from PKM2^apo^ and PKM2^FBP^ simulations, respectively, in addition to cluster C3 that was populated by sub-states from both simulations (**Fig. 4b**). The observed separation in the correlated motions revealed common conformational sub-states, suggesting that the preceding analysis of MD simulations of PKM2 captured FBP–dependent correlated motions.

We next subtracted the mutual information matrices identified in PKM2^apo^ from those in PKM2^FBP^ (**Fig. 4c**) to identify allosteric hub fragments (AlloHubFs) that are involved in the allosteric state transition. We found that the strength of the coupling signal between the AlloHubFs correlated with the positional entropy (**Fig. 4d**), corroborating the idea that local backbone flexibility contributed to the transmission of allosteric information. The top ten predicted AlloHubFs (named Hub_1_-Hub_10_) were spatially dispersed across the PKM2 structure including positions proximal to the A-A’ interface (Hub_5_ and Hub_6_), to the FBP binding pocket (Hub_9_), the C-C’ interface (Hub_10_), and within the B-domain (Hub_1_ and Hub_2_) (**Fig. 4e**). Remarkably, all AlloHubFs, with the exception of Hub_5_ and Hub_6_, coincided with minimal distance pathways^57^ between the FBP binding pocket and the active site (**Fig. 4e**). This observation further supported the hypothesis that the selected AlloHubFs propagate the allosteric effect of FBP.

### AlloHubF mutants disrupt FBP–induced activation of PKM2 or its sensitivity to Phe

We next generated *allosteric hub mutants* (AlloHubMs) (**Supplementary Fig. 8**) by substituting AlloHubF residues with amino acids that had chemically different side chains and were predicted to be tolerated at the respective position based on their occurrence in a multiple sequence alignment of 5381 pyruvate kinase orthologues (**Supplementary Fig. 9**). Among a total of 32 PKM2 AlloHubMs generated, we chose seven [I124G, F244V, K305Q, F307P, A327S, C358A, R489L (**Fig. 5a**)] for further experimental characterization because they expressed as soluble proteins and had very similar secondary structure to that of PKM2(WT), which suggested that the protein fold in these mutants was largely preserved (**Supplementary Fig. 10**). Importantly, the K_D_^FBP^ for all AlloHubMs were similar to PKM2(WT) (**Supplementary Fig. 11** and **Table 5**), with the exception of PKM2(R489L), which bound to FBP with low affinity.

**Figure 5:**
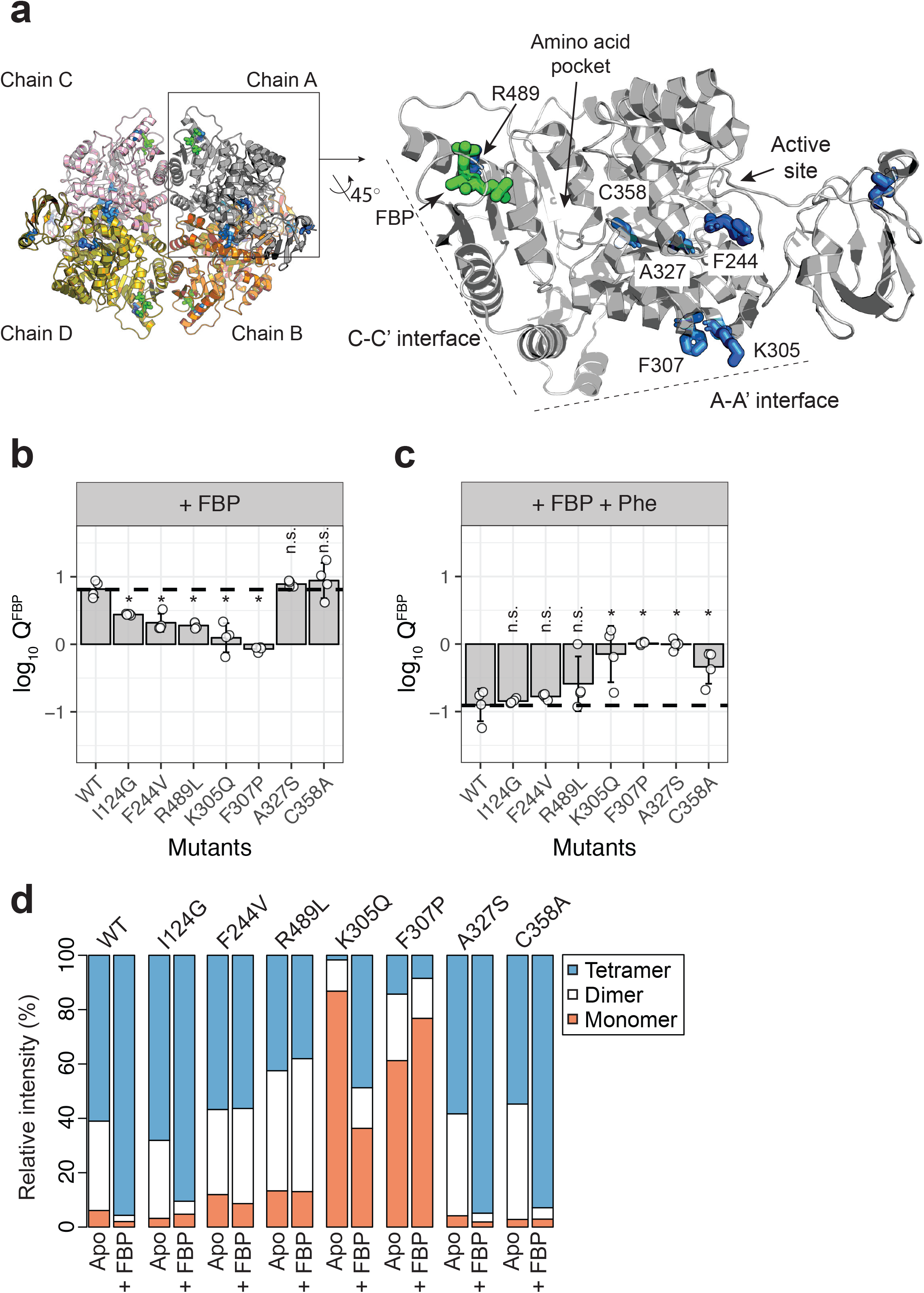
AlloHubF residue mutants interfere with FBP-induced PKM2 activation or its disruption by Phe. **(a)** AlloHubF mutants characterised in this study, shown on the PKM2 protomer structure. **(b)** The allosteric response of PKM2(WT) and AlloHubF mutant enzymatic activities to FBP, quantified by the allosteric coefficient Q, which denotes the change of the K_M_^PEP^ in the absence and in the presence of saturating concentrations of FBP (see Methods). A Q-coefficient > 0, indicates allosteric activation; and Q-coefficient < 0 indicates allosteric inhibition. The *Q*-coefficient for wild type PKM2 (WT) is shown as a dotted line for comparison. Each of the *Q*-coefficients of the AlloHubF mutants were statistically compared to PKM2(WT) using a Wilcoxon ranked-sum test (*n* = 4); a p-value < 0.05 was deemed significant (denoted by an asterisk); n.s.: not significant. **(c)** The magnitude of allosteric inhibition by Phe, in the presence of FBP, determined for PKM2(WT) and AlloHubF mutants, quantified by the allosteric co-efficient Q as in **(b)**. **(d)** Relative abundance of monomers, dimers and tetramers for PKM2 (WT) and PKM2 AlluHubF mutants in the absence or presence of saturating FBP.

**Table 5.**
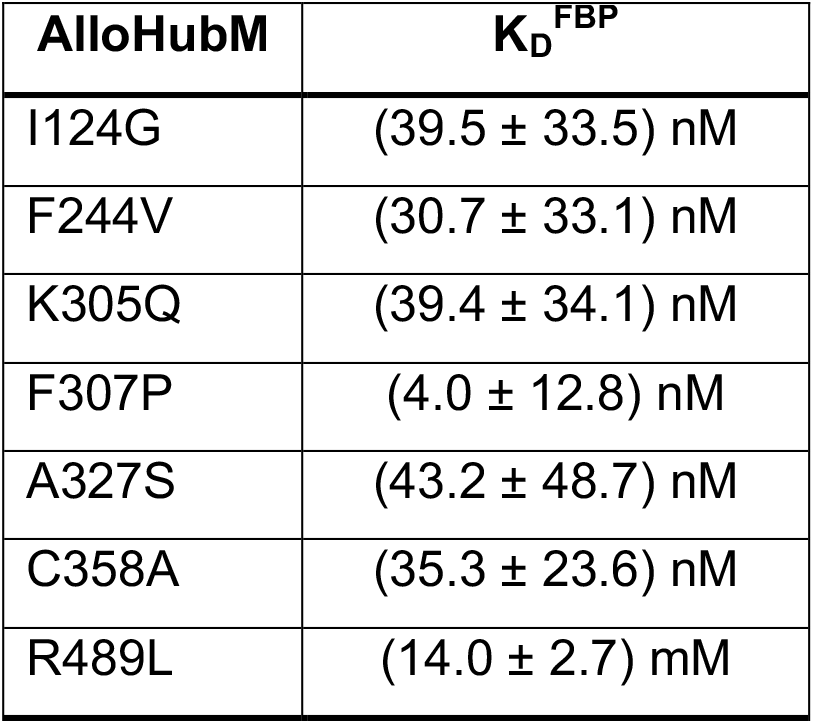
Apparent steady-state binding constant of FBP to the PKM2 allosteric hub mutants.

| **AlloHubM** | **K<sub>D</sub><sup>FBP</sup>** |
| --- | --- |
| I124G | (39.5 ± 33.5) nM |
| F244V | (30.7 ± 33.1) nM |
| K305Q | (39.4 ± 34.1) nM |
| F307P | (4.0 ± 12.8) nM |
| A327S | (43.2 ± 48.7) nM |
| C358A | (35.3 ± 23.6) nM |
| R489L | (14.0 ± 2.7) mM |

In order to quantify and compare the ability of FBP to activate AlloHubMs independently of varying basal activity, we determined the allosteric coupling constant^58,59^ log_10_Q^FBP^ defined as the log-ratio of the K_M_^PEP^ in the absence over the K_M_^PEP^ in the presence of FBP. PKM2 AlloHubMs I124G, F244V, K305Q, F307P and R489L showed attenuated activation by FBP (**Fig. 5b**) indicating that AlloHubMat successfully identified residues that mediate the allosteric effect of FBP. Similar to PKM2(WT), Phe addition significantly hindered FBP–induced activation in I124G, F244V, and R489L (**Fig. 5c**). In contrast, K305Q and F307P were allosterically inert, with no detectable response in activity upon addition of either FBP or Phe (**Fig. 5b,c**). While for PKM2(K305Q) this outcome could be explained by very low basal activity, PKM2(F307P) had a K_M_^PEP^ similar to that of PKM2(WT)^FBP^, indicating that this mutant displays a constitutively high substrate affinity (**Table 6**).

**Table 6.** Steady-state Michaelis-Menten kinetic parameters for AlloHubF mutants.

| AlloHubM | Ligand | $K_M^{\text{PEP}}$ (mM) | $k_{\text{cat}}$ ($\text{s}^{-1}$) | $k_{\text{cat}}/K_M^{\text{PEP}}$ ($\text{s}^{-1} \text{mM}^{-1}$) |
| --- | --- | --- | --- | --- |
| I124G | Apo | $1.07 \pm 0.13$ | $190.27 \pm 7.31$ | $177.82 \pm 56.23$ |
| | FBP | $0.29 \pm 0.04$ | $307.10 \pm 6.11$ | $1058.97 \pm 152.75$ |
| | FBP+Phe | $1.44 \pm 0.34$ | $221.92 \pm 17.26$ | $153.98 \pm 37.88$ |
| F244V | Apo | $1.14 \pm 0.11$ | $237.44 \pm 7.36$ | $210.44 \pm 27.15$ |
| | FBP | $0.55 \pm 0.07$ | $279.62 \pm 9.93$ | $522.42 \pm 87.51$ |
| | FBP+Phe | $1.15 \pm 0.30$ | $222.51 \pm 18.29$ | $195.87 \pm 55.46$ |
| K305Q | Apo | $0.01 \pm 0.01$ | $8.06 \pm 0.57$ | $790.01 \pm 605.56$ |
| | FBP | $0.01 \pm 0.04$ | $8.40 \pm 0.43$ | $1038.01 \pm 535.50$ |
| | FBP+Phe | $0.04 \pm 0.01$ | $12.21 \pm 1.40$ | $408.37 \pm 136.10$ |
| F307P | Apo | $0.13 \pm 0.01$ | $180.47 \pm 3.05$ | $1413.87 \pm 133.12$ |
| | FBP | $0.15 \pm 0.02$ | $227.20 \pm 5.02$ | $1508.53 \pm 190.71$ |
| | FBP+Phe | $0.21 \pm 0.03$ | $328.71 \pm 12.37$ | $1568.32 \pm 268.58$ |
| A327S | Apo | $1.37 \pm 0.42$ | $31.8 \pm 2.43$ | $23.21 \pm 5.79$ |
| | FBP | $0.17 \pm 0.02$ | $119.01 \pm 6.86$ | $700.06 \pm 343.00$ |
| | FBP+Phe | $0.15 \pm 0.02$ | $100.19 \pm 5.32$ | $667.93 \pm 266.00$ |
| C358A | Apo | $4.71 \pm 0.99$ | $214.73 \pm 21.40$ | $45.59 \pm 21.62$ |
| | FBP | $0.58 \pm 0.18$ | $191.13 \pm 16.22$ | $329.53 \pm 90.11$ |
| | FBP+Phe | $1.25 \pm 0.07$ | $199.01 \pm 21.87$ | $159.20 \pm 31.24$ |
| R489L | Apo | $0.69 \pm 0.19$ | $60.05 \pm 4.91$ | $89.38 \pm 36.93$ |
| | FBP | $0.36 \pm 0.10$ | $112.25 \pm 7.60$ | $317.20 \pm 125.71$ |
| | FBP+Phe | $1.44 \pm 0.40$ | $132.99 \pm 15.74$ | $158.19 \pm 122.56$ |

The mass spectra of the AlloHubMs (**Fig. 5d**) revealed a marked decrease in the intensity of tetramer and dimer peaks for PKM2(K305Q) and PKM2(F307P) compared to PKM2(WT), consistent with the position of these residues on the A-A’ interface, along which stable PKM2(WT) dimers are formed (**Supplementary Fig. 4a,b**). However, upon addition of FBP, F307P remained largely monomeric, whereas K305Q formed tetramers with a similar charge state distribution to PKM2(WT)^apo^ (**Supplementary Fig. 12a,b**). These observations, in addition to the varying ability of I124G, F244V, and R489L to tetramerise upon addition of FBP (I124G > F244V > R489L, **Supplementary Fig. 12a,b**), indicate that an impaired allosteric activation of these mutants by FBP cannot be accounted for by altered oligomerisation alone.

Intriguingly, two AlloHubMs, PKM2(A327S) and PKM2(C358A), retained intact activation by FBP (**Fig. 5b**), suggesting either that the amino acid substitutions were functionally neutral or that these residues are not required for FBP-induced activation. However, addition of Phe failed to attenuate FBP-induced activation of these AlloHubMs (**Fig. 5c**), indicating that residues A327 and C358 have a role in coupling the allosteric effect of Phe with that of FBP.

In summary, evaluation of the allosteric properties of AlloHubMs demonstrated that AlloHubMat successfully identified residues involved in FBP-induced allosteric activation of PKM2. Furthermore, this analysis revealed two residues that mediate a functional cross-talk between allosteric networks elicited from distinct ligand binding pockets on PKM2, thereby providing a mechanism by which distinct ligands synergistically control PKM2 activity.

## DISCUSSION

Allosteric activation of PKM2 by FBP is a prototypical and long-studied example of feedforward regulation in glycolysis^60^. However, PKM2 binds to many other ligands in addition to FBP, including inhibitory amino acids. It has been unclear, thus far, whether ligands that bind to distinct pockets elicit functionally independent allosteric pathways to control PKM2 activity or whether they synergise, and if so, which residues mediate such synergism. Furthermore, the role played by simultaneous binding of multiple ligands on the oligomerisation state of PKM2 remains elusive. Our work shows that FBP-induced dynamic coupling between distal residues functions, in part, to enable Phe to interfere with FBP-induced allostery. This finding points to a functional cross-talk between the allosteric mechanisms of these two ligands.

### AlloHubMat reveals residues that mediate a cross-talk between FBP- and Phe-induced allosteric regulation

Multiple lines of evidence suggested a functional cross-talk between the allosteric mechanisms of Phe and FBP. While Phe and FBP bind to spatially distinct pockets on PKM2, both ligands influence the mode of action of the other, without reciprocal effects on their binding affinities. Using native MS, we showed that, while FBP and Phe individually have opposing effects on PKM2 oligomerisation, they synergistically stabilise PKM2 tetramers. However, in contrast to PKM2^FBP^ tetramers, PKM2^FBP+Phe^ tetramers have low enzymatic activity. Conversely, the presence of FBP altered the kinetic mechanism of Phe inhibition: Phe alone did not decrease the *k_cat_*, but instead increased the K_M_^PEP^. With FBP present, Phe decreased both the apparent affinity for PEP and the maximal velocity of FBP-bound PKM2.

To identify residues that mediate the interaction between the Phe and FBP allosteric mechanisms, we analysed changes imposed by FBP binding on PKM2 dynamics. MD simulations of PKM2^apo^ and PKM2^FBP^, corroborated by IM-MS data, indicated that the protein does not undergo large conformational changes, in agreement with previously published small-angle X-ray scattering (SAXS) experiments that showed no FBP-driven change in the radius of gyration of PKM2 tetramers^36^. This suggested that ligand-induced conformational changes are likely limited to subtle backbone re-arrangements and side-chain motions. Based on our previous approach^61–63^, we calculated the mutual information between sampled conformational states from MD trajectories encoded in a coarse-grained representation within the framework of a structural alphabet. Nevertheless, given that proteins have been shown experimentally^55,64–66^ and computationally^67,68^ to sample multiple conformational sub-states with distinct structural properties, explicitly identifying allosteric signals that are representative of the ensemble of protein sub-states is crucial. In order to derive the network of correlated motions from multiple MD trajectories and obtain an ensemble-averaged mutual information network, we developed AlloHubMat. AlloHubMat overcomes some of the limitations^56,64^ of previous approaches, and predicts networks of allosteric residues from MD simulations using a consistent numerical framework to measure time-dependent correlated motions. This approach enables both the extraction of consensus allosteric networks from replicate simulations of a protein in a given liganded state, and also the comparison of such consensus networks to each other.

AlloHubMat revealed candidate residues involved in the allosteric effect of FBP on PKM2 activity. Mutagenesis at several of these positions (I124G, F244V, K305Q, F307P, R489L) disrupted FBP–induced activation demonstrating that AlloHubMat successfully identified *bona fide* mediators of FBP allostery. Some AlloHub mutations also disrupt Phe-induced inhibition suggesting that FBP and Phe elicit allosteric effects through partially overlapping networks of residues. In contrast, PKM2(F244V) specifically disrupted FBP–induced activation, while maintaining the propensity for allosteric inhibition by Phe. Conversely, mutation of A327 and C358 preserved the ability of FBP to regulate PKM2 but prevented the inhibitory effect of Phe on PKM2^FBP^ enzymatic activity. This finding indicated a role for A327 and C358 in mediating the cross-talk between the allosteric mechanisms elicited by Phe and FBP that allows the former to interfere with the action of the latter. Notably, identification of C358 as an allosteric hub could explain why a chemical modification at this position perturbs PKM2 activity by oxidation^13^. None of the characterised mutants fall within positions 389-429 that differ between PKM2 and the constitutively active PKM1, suggesting that residues that confer differences in the allosteric properties of these two isoforms are dispersed throughout the protein. Our findings highlight the importance of experimentally evaluating the functional role of allosteric residues predicted by computational methods not only in the context of the allosteric effector under investigation but also in response to other potential allosteric effectors that may also be unidentified.

Interestingly, Zhong *et al*.^69^ recently reported that adenosine monophosphate (AMP) and glucose-6-phosphate (G6P) synergistically activate *M. tuberculosis* pyruvate kinase. While the binding of AMP occurs at a pocket equivalent to that of PKM2 for FBP, G6P binds to a different pocket that is also distinct from the equivalent amino acid interaction site on PKM2, indicating an additional allosteric integration mechanism that is similar to the one we describe here. It is therefore tempting to speculate that allosteric synergism upon concurrent binding of different ligands may occur more commonly that previously appreciated.

### AlloHubMs reveal insights into the relationship between PKM2 oligomerisation and enzymatic activity

Changes in oligomerisation have been intimately linked to the regulation of PKM2 activity. AlloHuMs A327S and C358A retained both their ability to tetramerise and increase their activity in response to FBP. Furthermore, FBP fails to shift the oligomer equilibrium and does not activate AlloHuMs F307P and R489L. In this context, we found that Phe inhibited the activity of PKM2^apo^, with concomitant loss of tetramers, however, Phe inhibition of PKM2^FBP^ occurred within the tetrameric state. The mechanism by which Phe regulates PKM2 oligomerisation is controversial. Our finding that Phe destabilises tetramers is in agreement with previous studies^38^, but contrasts with recent reports that Phe stabilises a low activity T-state tetramer^29,31^. Critically, we find that Phe and FBP synergistically promote PKM2 tetramerisation, raising the possibility that the mode of Phe action described in Morgan *et al*. and Yuan *et al*.^31^ is confounded by the presence of *residual* FBP. It is unclear whether partial FBP occupancy is accounted for in these studies, as the FBP saturation status of PKM2 is not detailed. Co-purification of FBP with recombinant PKM2 has been previously observed^29,35,46^ and in purifying recombinant PKM2 for our study, we detected up to more than 0.75 fractional saturation with co-purified FBP. Furthermore, initial attempts, by others, to crystallise Phe-bound PKM2 without FBP required mutation in the protein (R489L) to abrogate FBP binding^29^, although structures of PKM2(WT) bound to Phe have now been obtained^31^. Based on our findings, we speculate that the stabilisation of PKM2 tetramers by Phe observed by Morgan *et al*.^29^ and Yuan *et al*.^31^ could be attributed to significant amounts of residually-bound FBP co-purifying from *E. coli*. Consistent with this hypothesis, the small shift in the conformational arrangement (^DT^CCS_He_) of PKM2^FBP^ tetramers upon addition of Phe (PKM2^FBP+Phe^) may be indicative of subtle conformational changes that reflect a transition from an active R-state to an inactive T-state described before^29^. In support of this interpretation, the ^DT^CCS_He_ of PKM2^FBP+Phe^ tetramers closely resembles that of PKM2^apo^.

Therefore, beyond its immediate goals, our study has broader implications for understanding the regulation of PKM2 as it suggests that activation can be uncoupled from tetramerisation. This idea resonates well with findings from MD simulations indicating that dynamic coupling between distal sites upon FBP binding can occur in the PKM2 protomer, and suggest that allostery is encoded in the protomer structure^70–72^. Furthermore, a patient-derived PKM2 mutant (G415R) occurs as a dimer that can bind to FBP but cannot be activated and does not tetramerise^36^. Moreover, SAICAR can activate PKM2(G415R) dimers, in the absence of tetramerisation^36^. The finding that enzymatic activation is not obligatorily linked to tetramerisation is important for studies in intact cells, where distinction between the T- and R-states is not possible and therefore the oligomeric state of PKM2 is frequently used to infer activity^13,14,18,20,22,42–44^.

### Multiple allosteric inputs in the context of intracellular concentrations of allosteric effectors and other modifications

Our findings also highlight the importance of interpreting allosteric effects detected *in vitro* in the context of intracellular concentrations of the respective effectors. Reversible binding of FBP to PKM2 *in vitro* is a well-studied regulatory mechanism. However, our findings reveal that FBP concentrations far exceed the concentration needed for full saturation of PKM2 and, under steady-state growth conditions, a significant fraction of PKM2 is already bound to the activator FBP, even in the context of other regulatory cues, such as PTMs, that may influence ligand binding. Furthermore, our results showed that PKM2 inhibition by Phe can occur even under conditions of saturating FBP, both with purified PKM2 and in cell lysates. Taken together, these observations indicate that inhibition by Phe constitutes a physiologically relevant regulatory mechanism that may contribute to maintaining PKM2 at a low activity state, as is often found in cancer cells^14^.

Intriguingly, other amino acids can bind the same pocket as Phe, including activators Ser^25^ and His^31^. Therefore, it is likely that amino acids combined, rather than individually, control PKM2, as also supported by recent findings by Yuan *et al*.^31^. Further work is warranted to understand how all of these cues are integrated by PKM2.

In summary, our findings reveal that allosteric inputs from distinct ligands are integrated to control the enzymatic activity of PKM2. This is analogous to multiple-input-single-output (MISO) controllers in control system engineering in which multiple transmission signals (allosteric ligands) are integrated to a single receiving signal (enzyme activity)^73^. It is likely that many proteins can bind to multiple allosteric ligands that co-exist in cells. Whether a systems-control ability in integrating the allosteric effects of multiple ligands with opposing functional signals is a general property of other proteins is not known. Identification of mutations, using AlloHubMat, that perturb allosteric responses to specific ligands, alone or in combination, provides an essential means to study both the mechanistic basis of allosteric signal integration as well as the functional consequences of combinatorial allosteric inputs on cellular regulation.

## ACKNOWLEDGMENTS

We would like to thank all members of the Anastasiou and Fraternali laboratories for fruitful discussions, feedback and technical advice. We thank Mariana Silva dos Santos and James MacRae (Crick Metabolomics Science Technology Platform) for discussions and advice on metabolite measurements, Nicola O’Reilly (Crick Peptide Chemistry Science Technology Platform) for the synthesis of labelled peptides, and Colin Davis (Crick Proteomics Science Technology Platform) for technical assistance with proteomics. We thank Jakub Ujma and Waters Corp. for the use of the prototype SID instrument and the Wysocki Group (Ohio State University) for helpful discussions on SID tuning. AT is supported by BBSRC grants BB/L002655/1, BB/L016486/1 which is a studentship with additional financial support from Waters Corp. This work was funded by the MRC (MC_UP_1202/1) and by the Francis Crick Institute, which receives its core funding from Cancer Research UK, the UK Medical Research Council and the Wellcome Trust to DA (FC001033).

## AUTHOR CONTRIBUTIONS

JAM designed, performed and analysed all experiments, wrote all the software and wrote the manuscript. AT performed nESI-MS and IM-MS analyses. LM, SM and PCD assisted with biophysical experiments. LF performed metabolite concentration measurements. VE and APS performed proteomics experiments. JK assisted with software development, provided guidance and supervision on computational aspects of the study. PEB supervised and advised on nESI-MS and IM-MS experiments. FF conceived and supervised computational aspects of the study. DA conceived and supervised the study, designed experiments, interpreted data and wrote the manuscript. All authors read and commented on the manuscript.

## SUPPLEMENTARY FIGURE LEGENDS

**Supplementary Figure 1:**
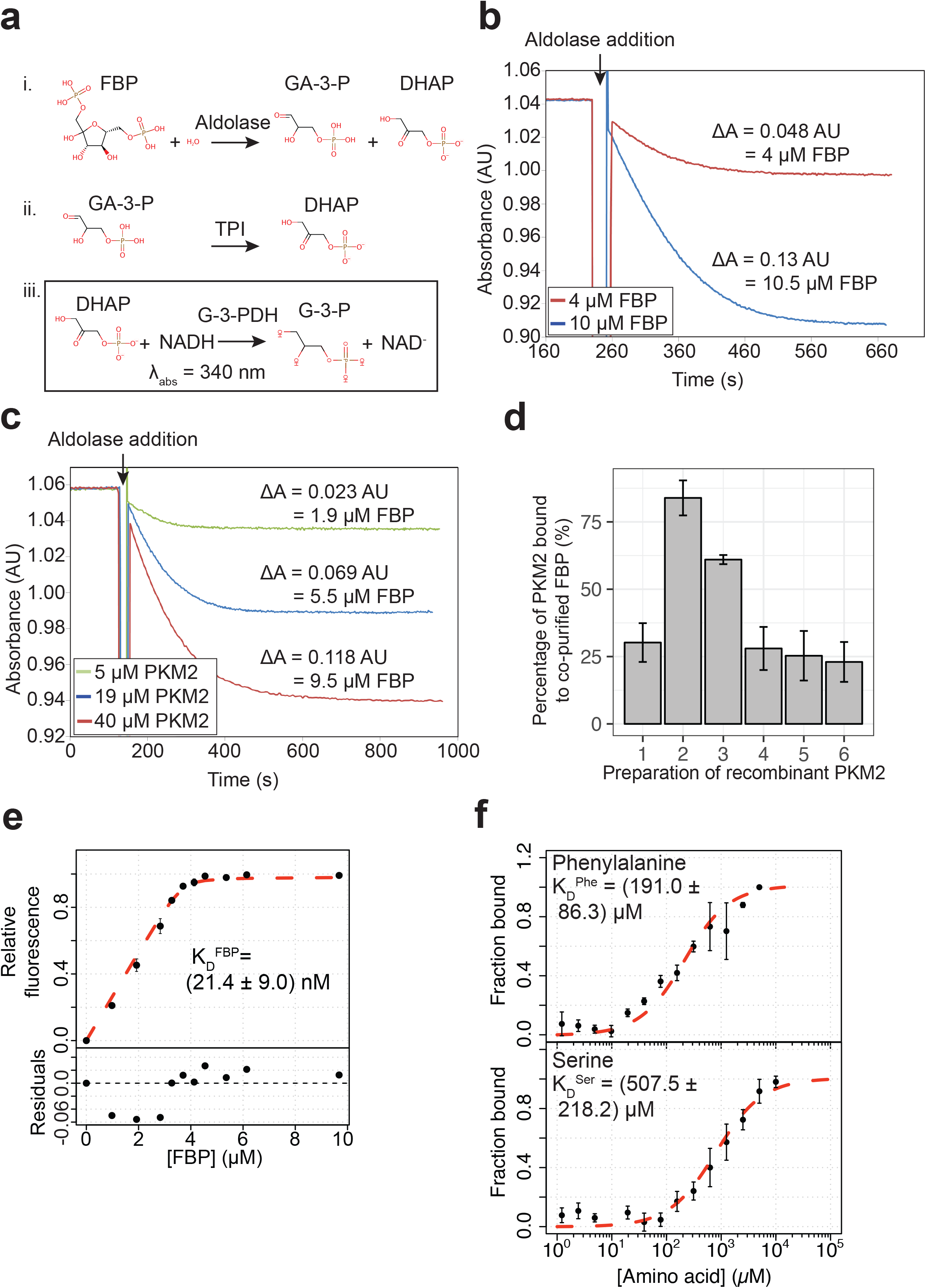
FBP binds to PKM2 with nM affinity. **(a)** FBP co-purified with recombinant PKM2 was quantified using a spectrophotometric coupled-enzyme assay that employs aldolase, triose phosphate isomerase (TPI) and glycerol-3-phosphate dehydrogenase (G-3-PDH). The final step of the three-enzyme reaction involves the oxidation of NADH, leading to a decrease in absorbance at 340 nm. **(b)** The aldolase coupled assay in **(a)** can reliably detect as low as 4 μM FBP. Sensitivity of the aldolase assay was tested by adding 4 or 10 μM purified FBP, and oxidation of NADH was initiated by adding the aldolase enzyme following an equilibration period. Amount of FBP was calculated from the measured decrease in the amount of NADH (one molecule of FBP consumed for every two molecules of NADH oxidised). **(c)** Levels of FBP detected in increasing amounts of recombinant PKM2. Indicated amounts of purified PKM2 were heat-precipitated at 95 °C to release any co-purified FBP and the supernatant was used for FBP quantification with the aldolase assay as in **(a)**. **(d)** Amounts of FBP quantified, as in **(a)**, for six independent preparations of purified recombinant PKM2. The amounts of co-purified FBP are represented as a proportion of the amount of recombinant PKM2. Preparations of PKM2 determined to have < 25% co-purifying FBP were used throughout this study. **(e)** Affinity of D-fructose-1,6-bisphosphate (FBP) for PKM2, determined by fluorescence emission spectroscopy (*λ*_EX_ = 280 nm, *λ*_EM_ = 290 – 450 nm) measuring fluorescence of PKM2 tryptophan residues, two of which are proximal to the FBP binding pocket. Fluorescence emission intensities at 325 nm and 350 nm were recorded with increasing concentrations of added ligand. The apparent binding constant (K_D_^FBP^) was estimated from a 1:1 binding model, fit to the experimental data using a non-linear least-squared fitting procedure (see Methods). The averages and standard deviations of four independent experiments are shown. **(f)** Affinity of Phe and Ser for PKM2 determined using microscale thermophoresis (MST) measurements of fluorescein-labelled PKM2 (30 nM). Indicated binding affinities were estimated using a 1:1 binding model. The average and standard deviations of four independent experiments are shown.

**Supplementary Figure 2:**
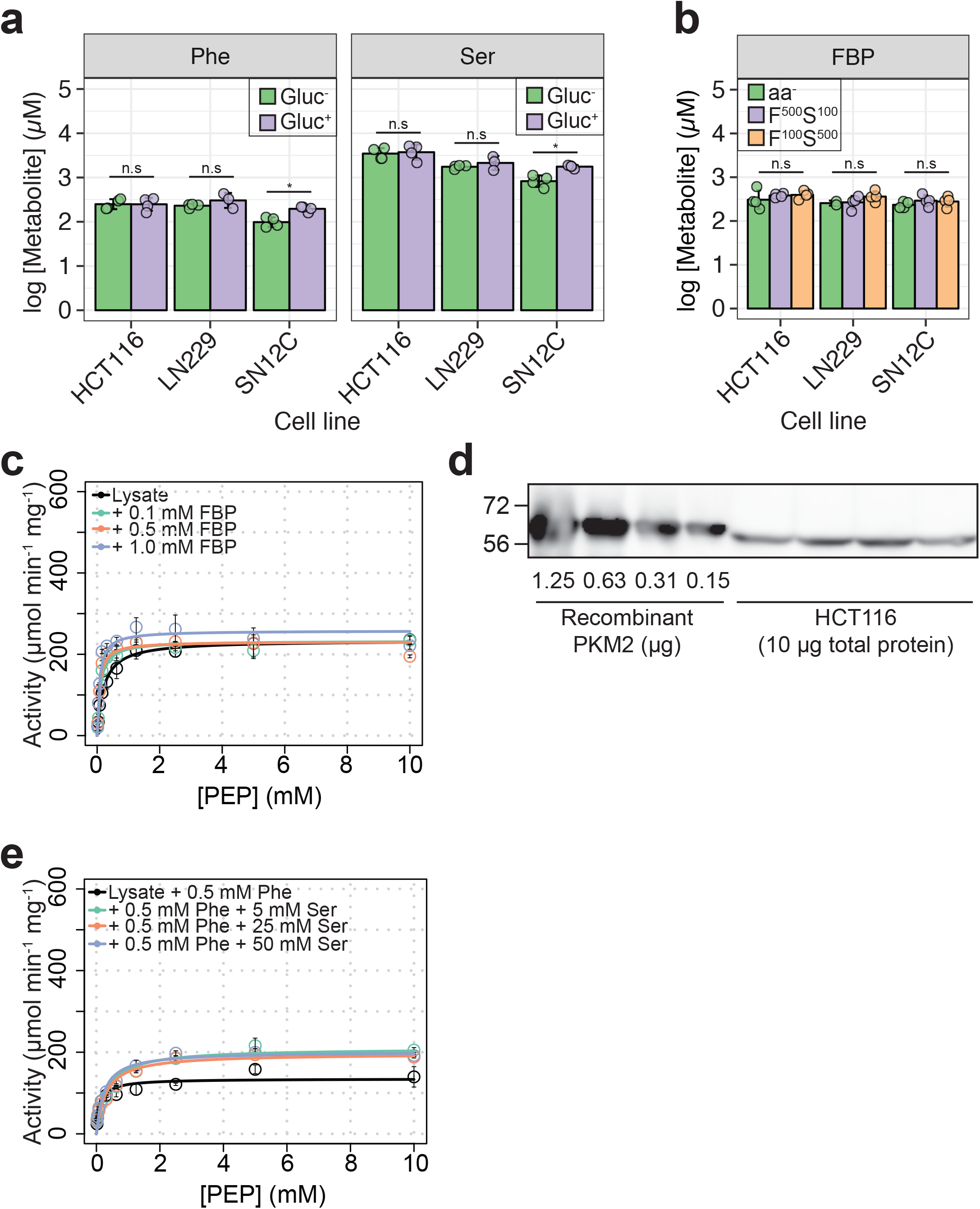
Metabolite levels and PKM2 activity in cells. **(a)** [Phe]_ic_ and [Ser]_ic_ in cells as in **Fig. 1a** showing no significant changes upon changes of glucose concentration in media. Statistical significance determined as in **Fig. 1a**. **(b)** [FBP]_ic_ in cells as in **Fig. 1c** showing no significant changes depending on Ser or Phe concentration in HBSS. Statistical significance determined as in **Fig. 1a. (c)** PKM2 activity as in **Fig. 1f** but in lysates of HCT116 cells cultured in RPMI without glucose (Gluc^−^). **(d)** Quantification of PKM2 in HCT116 lysates. Western blot of recombinant PKM2 or HCT116 cell lysates (Gluc+ media) was probed with a PKM2 antibody. An interpolation of the linear fit of the band intensities of four amounts of recombinant PKM2 was used to estimate the amounts of PKM2 protein per 1 μg of total protein in HCT116 cell lysates, in order to determine specific activity in panels **(c)** and **(e)**, and in **Fig. 1f and 1g. (e)** PKM2 activity in HCT116 cell lysates as in **Fig. 1g** but with 0.5 mM Phe and increasing amounts of Ser, as indicated.

**Supplementary Figure 3:**
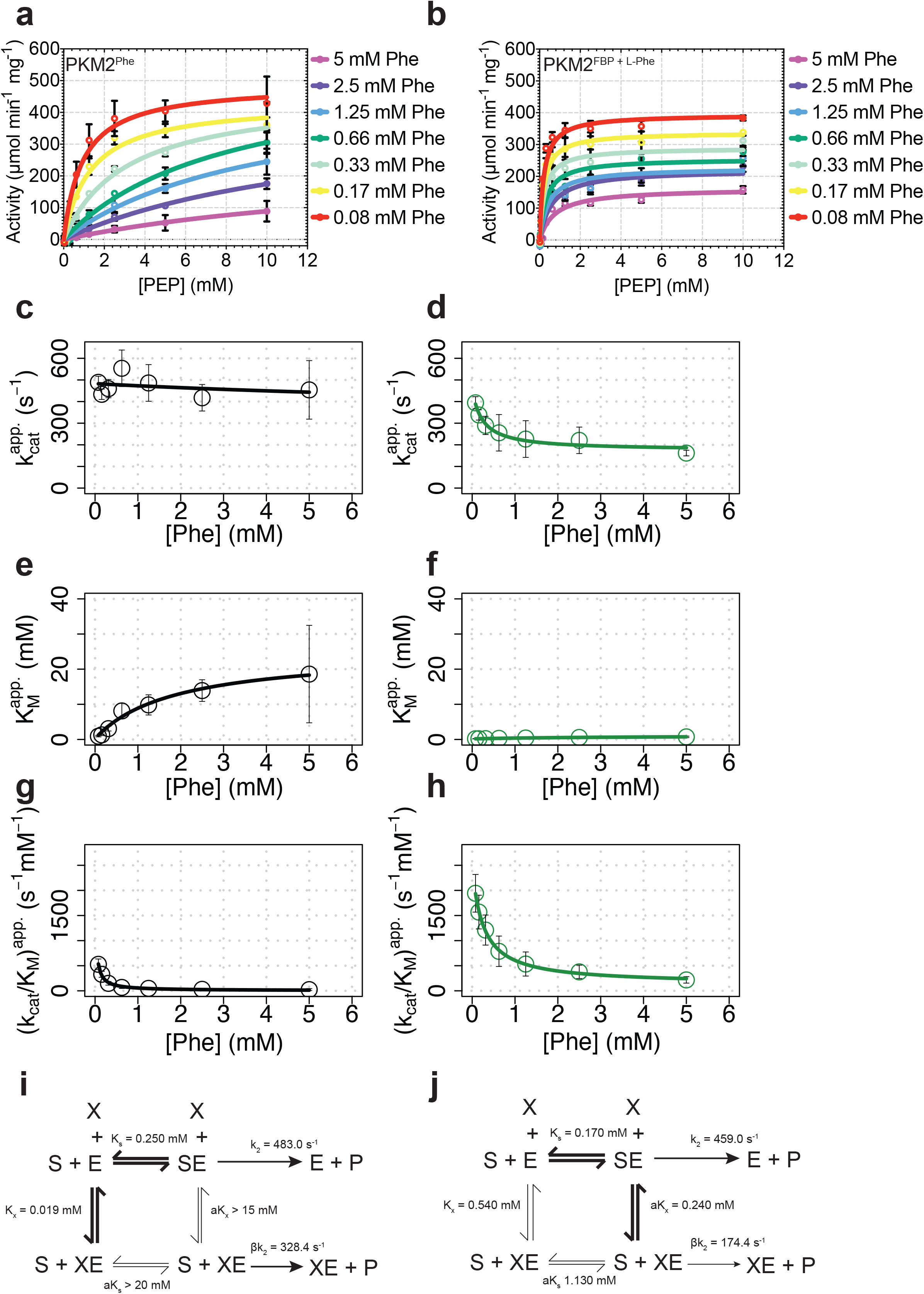
FBP influences the kinetics of PKM2 inhibition by Phe. **(a)** Purified recombinant PKM2 activity was assayed over a range of phosphoenolpyruvate (PEP) concentrations and with the indicated concentrations of phenylalanine (Phe). The averages and standard deviations of four separate experiments are shown. Initial velocity curves were fitted using the Michaelis-Menten model. **(b)** PKM2 activity was assayed in the presence of increasing concentrations of the inhibitor phenylalanine, as in **(a)**, in the presence of a constant concentration of 2 μM FBP. Rate curves were fitted to Michaelis-Menten kinetic models, from which the parameters k_cat_, K_M_^PEP^ and k_cat_/K_M_^PEP^ were computed for Phe inhibition of PKM2^apo^ (black; **c, e** and **g**) and PKM2^FBP^ (green; **d, f** and **h**). The dependence of the three steady-state kinetic parameters k_cat_, K_M_^PEP^ and k_cat_/K_M_^PEP^ were fit to the steady-state *modifier-rate* equations (see Methods). **(i)** The steady-state mechanism of allosteric inhibition is shown from a single-substrate-single-modifier representation of PKM2 catalysis; where *E* is the enzyme, *S* is the substrate PEP and *X* is the allosteric effector Phe. Equilibrium constants were assigned from fittings of the dependence of the kinetic parameters on the concentration of Phe, determined above. The mechanism of PKM2 inhibition by Phe in the absence of FBP is found to be hyperbolic-specific^51^. **(j)** The mechanism of Phe inhibition of PKM2^FBP^ was determined as in **(i)**, and was found to be of the variety of hyperbolic-mixed inhibition^51^.

**Supplementary Figure 4:**
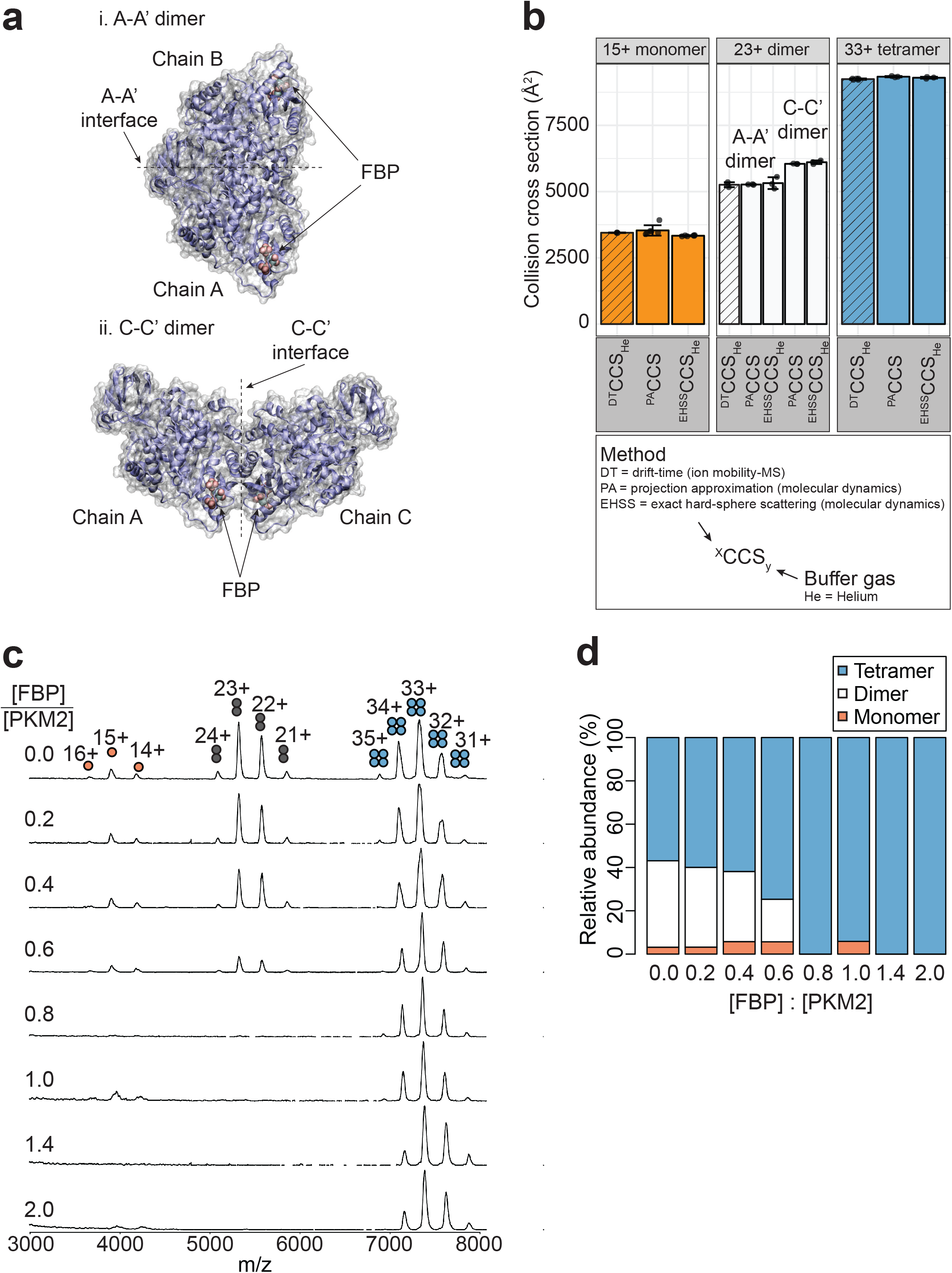
FBP promotes a dose-dependent conversion of PKM2 monomers and A-A’ dimers into the tetrameric species. **(a)** Representation of the two possible dimer assemblies based on PKM2 symmetry: i. the A-A’ dimer (protomers associated along the A-A’ interface) and ii. the C-C’ dimer (protomers associated along the C-C’ interface). **(b)** Evidence from IM-MS that the observed PKM2 dimeric species adopt the A-A’ configuration. The collisional cross section (^DT^CCS_He_) of 15+ monomer species (orange), 23+ dimer species (white) and 33+ tetramer species (blue) PKM2^apo^ were experimentally determined using ion mobility-mass spectrometry (IM-MS; striped bars). The exact hard sphere collision cross section (^EHSS^CCS_He_) and projection approximation collision cross section (^PA^CCS) were calculated from short *in vacuo* molecular dynamics simulations of 15+ monomeric (orange), 23+ dimer (white) and 33+ tetrameric (blue) PKM2^apo^. The ^DT^CCS_He_ of the dimeric species is very similar to that of the A-A’ dimer calculated by both ^EHSS^CCS_He_ and ^PA^CCS. **(c)** FBP-dose-dependent conversion of PKM2 monomers and dimers into tetramers. Mass spectra of PKM2 obtained following pre-incubation with increasing amounts of FBP resulting in the indicated FBP/PKM2 ratios. **(d)** Relative abundances of PKM2 monomers, dimers and tetramers determined from **(c)**.

**Supplementary Figure 5:**
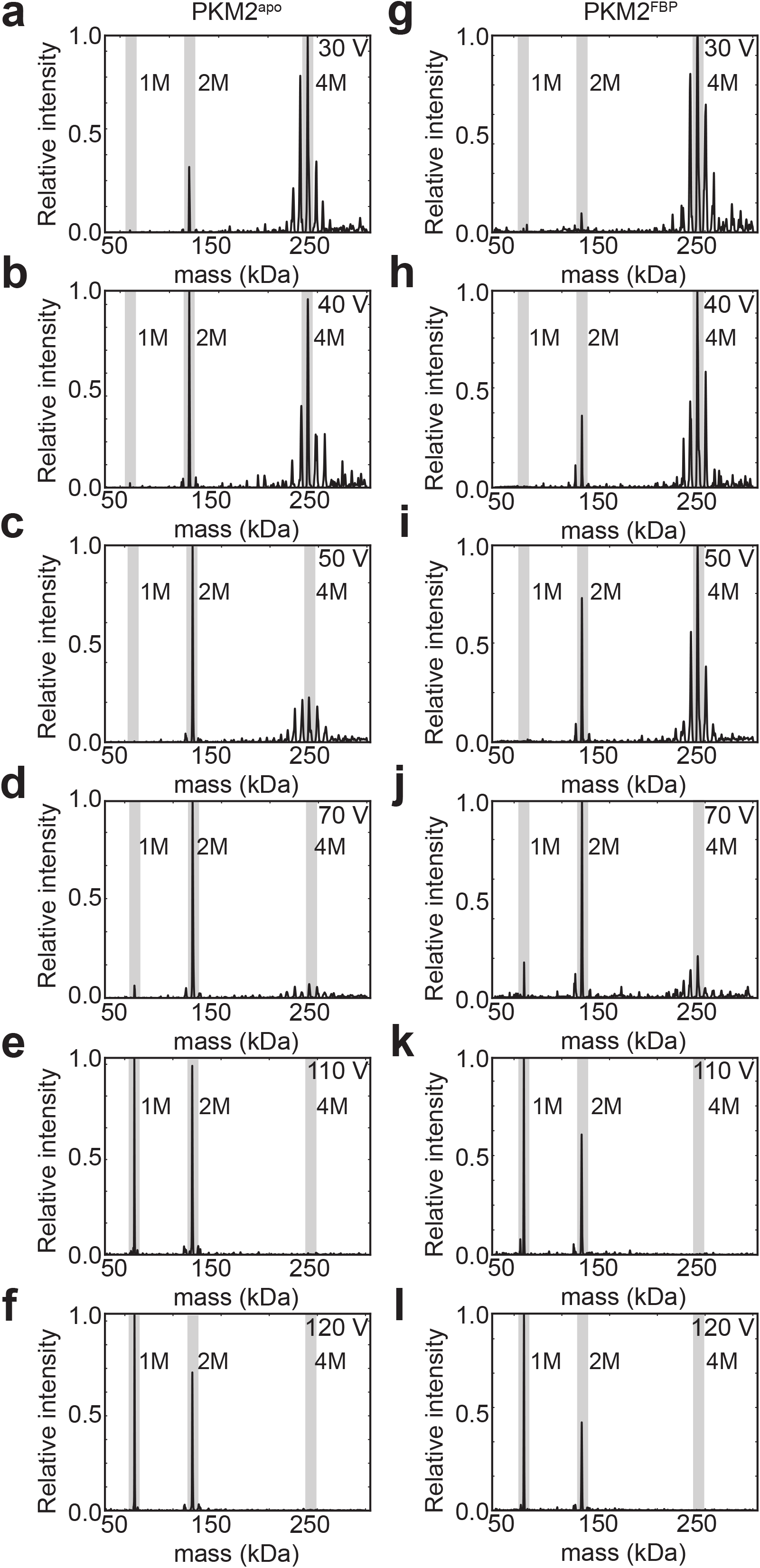
FBP binding increases the stability of PKM2 tetramers. Mass de-convolved spectra showing the relative abundance of oligomeric species produced from the 33+ charge state precursor of tetrameric PKM2 at increasing collision voltages (40 V – 120 V; lab-frame), obtained using surface-induced dissociation (SID) (see Methods) for PKM2^apo^ **(a-f)** and PKM2^FBP^ **(g-l)**. The positions corresponding to the tetramer, dimer and monomer species are highlighted in grey.

**Supplementary Figure 6:**
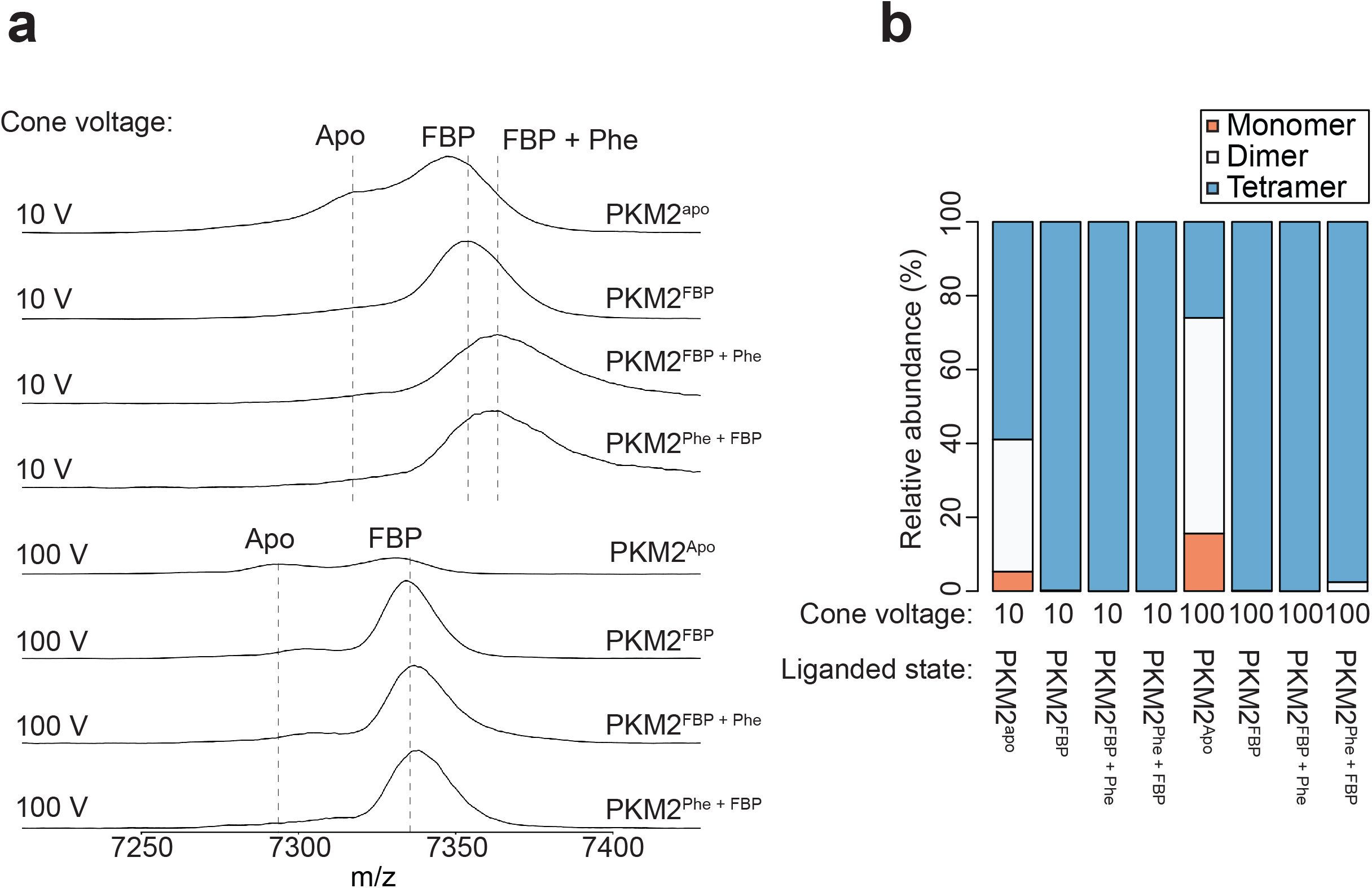
Phe and FBP can simultaneously bind to PKM2. **(a)** Mass spectra of PKM2 were acquired in the absence of any added ligands (PKM2^apo^), and following addition of stoichiometric amounts of FBP (PKM2^FBP^); addition of 400 μM Phe to 10 μM PKM2 pre-incubated with 10μM FBP (PKM2^FBP + Phe^); or addition of 10 μM FBP to 10 μM PKM2 pre-incubated with 400 μM Phe (pKM2^Phe + FBP^). Spectra were acquired with the cone-voltage of the electrospray-ionisation (ESI) source set at 10 V. An *m/z* shift is observed upon the addition of FBP, and a further shift is observed upon addition of Phe after FBP binding. The peaks are collisionally converted to an FBP-like state by increasing the cone voltage of the ESI source to 100 V, resulting in removal of phenylalanine. Positions of the m/z peaks for PKM2^apo^, PKM2^FBP^, PKM2^FBP + Phe^ and PKM2^Phe + FBP^ are shown as dashed lines at each of the two cone voltage conditions for reference. **(b)** The relative abundances of monomer (orange) dimer (white) and tetramer (blue) species were quantified from the spectra in **(a)**.

**Supplementary Figure 7:**
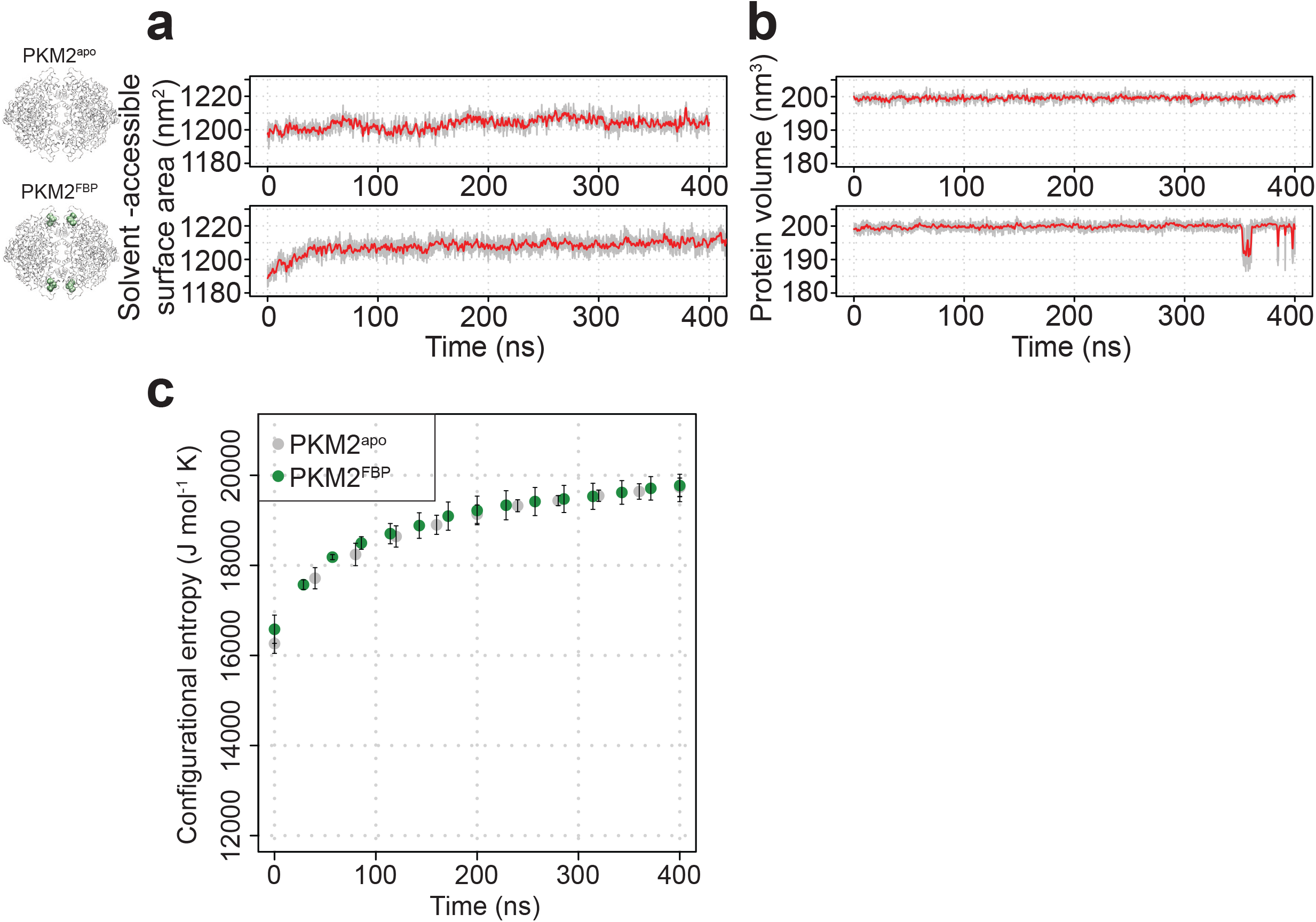
Analyses of MD simulations of PKM2. **(a)** Five separate molecular dynamics (MD) simulations of PKM2^apo^ and PKM2^FBP^ analysed for solvent-accessibility. The grey trace shows the variance of the solvent-accessibility over the replicate simulations and the red trace is the running average of the variance. **(b)** MD simulations as in **(a)** but analysed for volume. **(c)** The time-resolved configurational entropy was calculated for each of the MD simulations of PKM2^apo^ and PKM2^FBP^ in **(a, b)** using the Schlitter formula (see Methods). Mean values and standard deviations are shown for each of the trajectories.

**Supplementary Figure 8:**
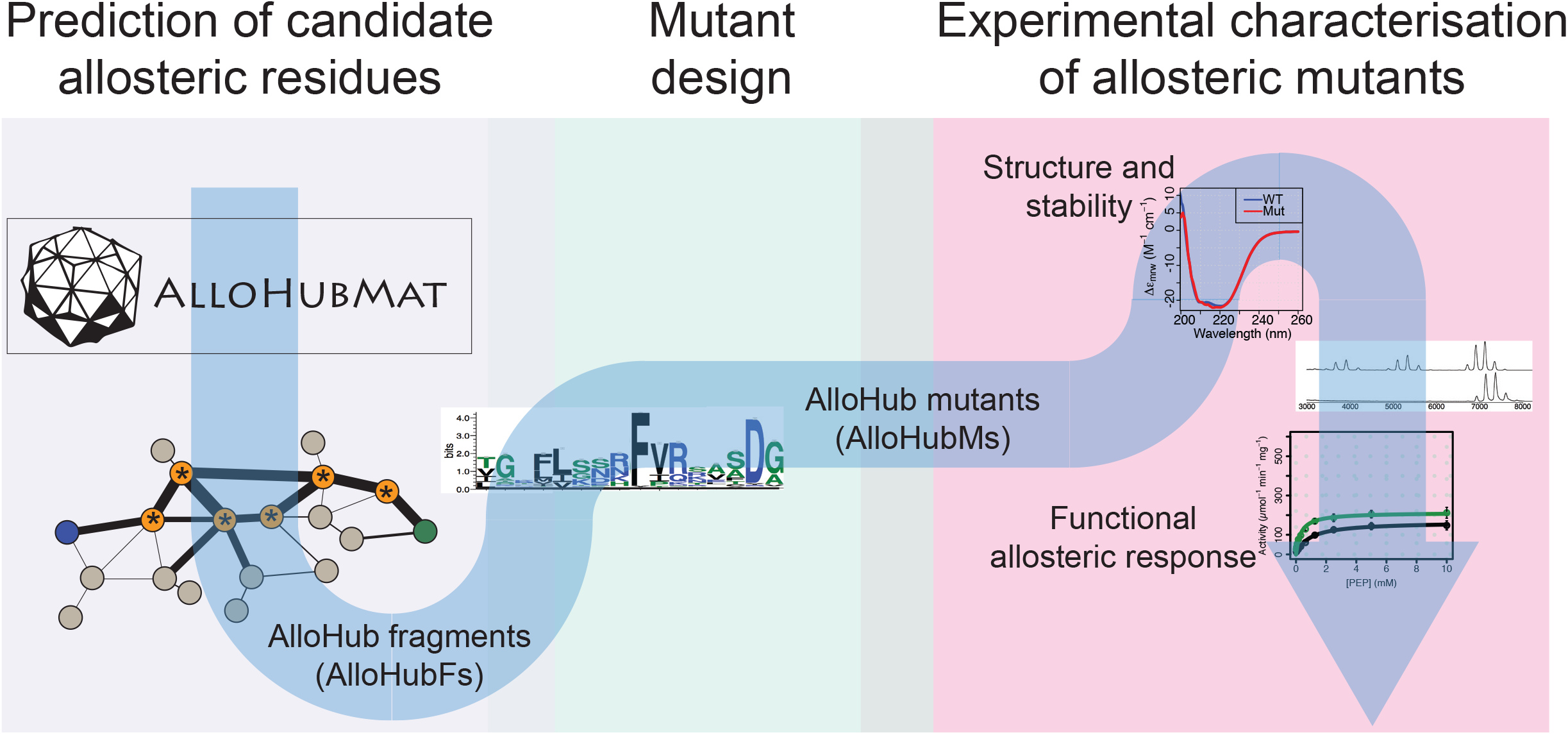
Schematic depicting the integrated computational and experimental strategy used to identify protein residues involved in allosteric regulation. Molecular dynamics (MD) simulations were analysed using the AlloHubMat analysis software (**Fig. 4a**) to generate a network of correlated protein dynamics that was used to identify allosteric hub fragments (AlloHubFs). The design of mutant variants (AlloHubMs) was guided by protein sequence conservation analysis of residues comprising AlloHubFs. AlloHubMs were experimentally characterised for their structural integrity using CD spectroscopy, for their oligomerisation state using ESI-MS, and enzymatic properties to assess their allosteric response to FBP and the ability of Phe to interfere with it. See text and methods for details.

**Supplementary Figure 9:**
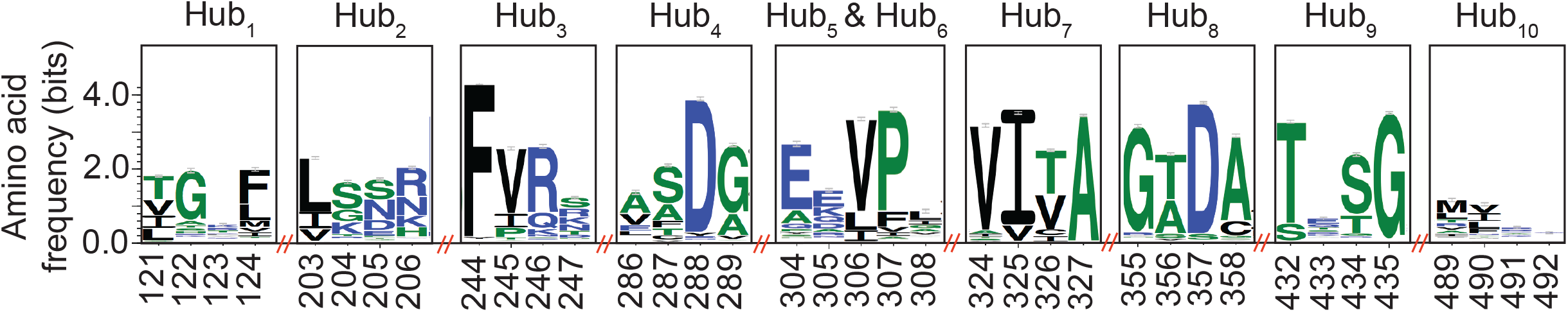
Sequence conservation analysis of AlloHubFs. A multiple sequence alignment of PKM2 homologues was generated through a BLAST search of the NCBI non-redundant protein database^75^. Sequences with a pairwise identity of greater than 0.8 were removed, leaving a total of 5381 homologues, which were aligned using the HMMER suite^76^. Sequence conservation logo plots were generated from the alignment for each of the top-ten predicted AlloHubF residues.

**Supplementary Figure 10:**
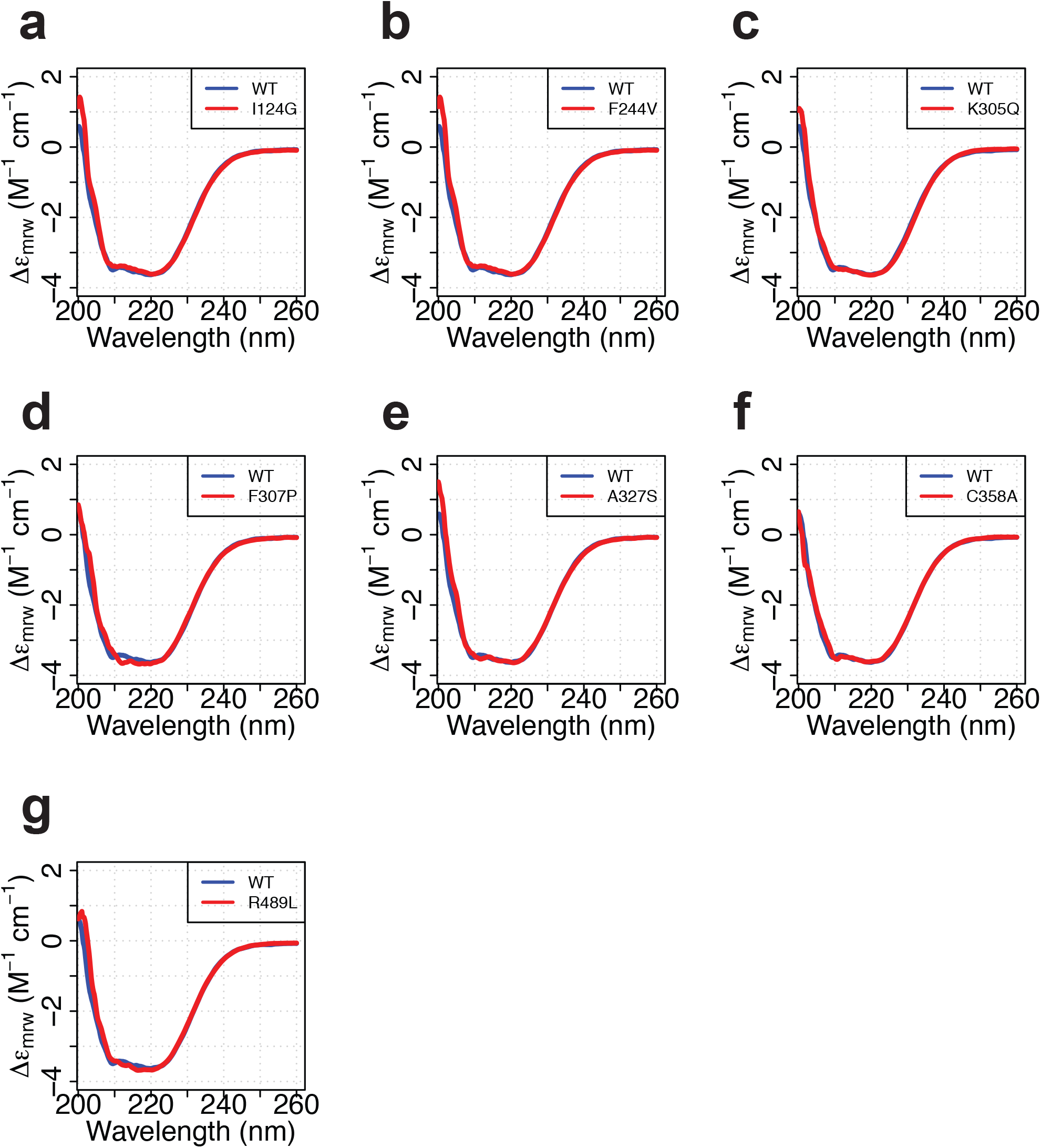
Purified AlloHubF mutants have similar secondary structure content to PKM2(WT). **(a-g)** Smoothened far-UV circular dichroism spectra (200 – 260 nm) of each of the AlloHubMs (red) superimposed to that of PKM2(WT) (blue).

**Supplementary Figure 11:**
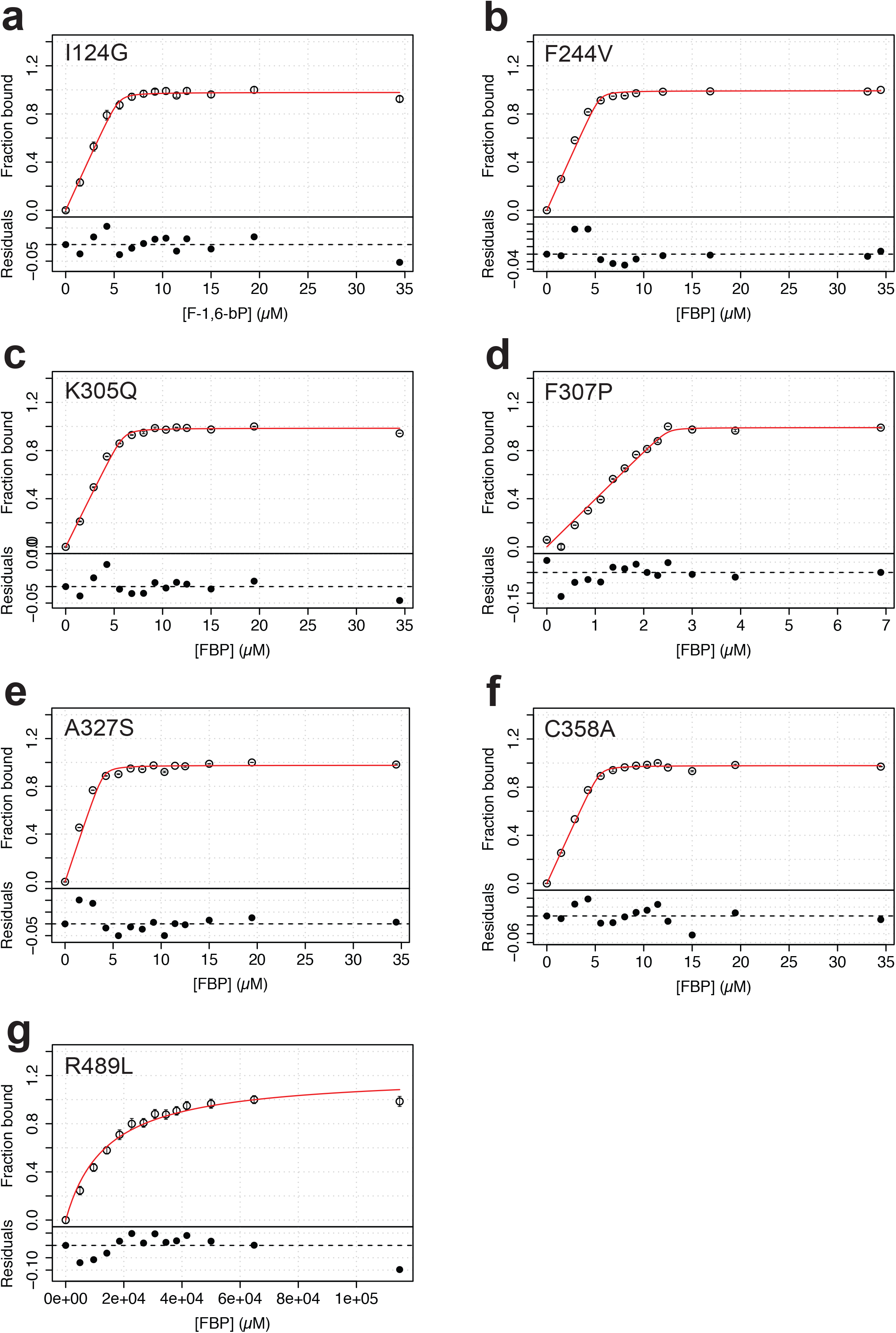
FBP has a similar affinity for AlloHubMs to that for PKM2(WT), with the exception of PKM2(R489L). **(a-g)** Fluorescence emission spectroscopy measurements were used to monitor binding of FBP to PKM2 for each of the AlloHubF mutants. Each binding curve was fit to a 1:1 binding model (see Methods), shown as a red line.

**Supplementary Figure 12:**
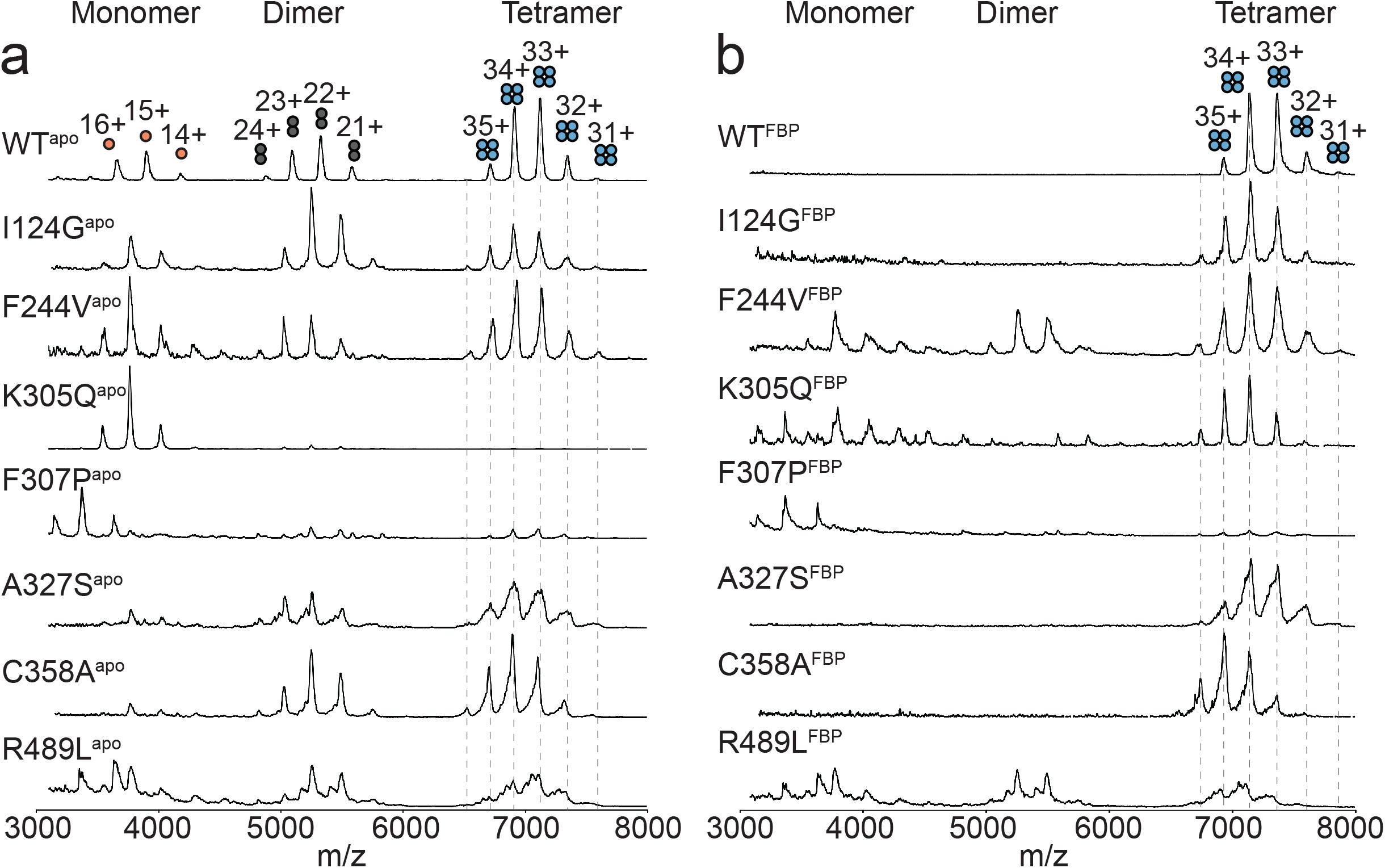
Native mass spectra of the AlloHubMs. **(a)** nESI-MS mass spectra of the AlloHubMs in the absence of any added ligands. **(b)** As in (a) but in the presence of FBP at a protein-to-ligand molar ratio of 1:10.

## METHODS

### Metabolite analyses by liquid chromatography-mass spectrometry (LC-MS)

The LC-MS method was adapted from Zhang *et al*. (2012)^77^. Briefly, samples were injected into a Dionex UltiMate LC system (Thermo Scientific; Waltham MA, USA) with a ZIC-pHILIC (150 mm × 4.6 mm, 5 μm particle) column (Merck Sequant; MilliporeSigma, Burlington MA, USA). A 15 minute elution gradient of 80% Solvent A to 20% Solvent B was used, followed by a 5 minute wash of 95:5 Solvent A to Solvent B and 5 min re-equilibration, where Solvent B was acetonitrile (Optima HPLC grade; Sigma Aldrich, St. Louis MS, USA) and Solvent A was 20 mM ammonium carbonate in water (Optima HPLC grade; Sigma Aldrich, St. Louis MS, USA). Other parameters were used as follows: injection volume 10 μL; autosampler temperature 4 °C; flow rate 300 μL/min; column temperature 25 °C. MS was performed using positive/negative polarity switching using an Q Exactive Orbitrap (Thermo Scientific; Waltham MA, USA) with a HeSI II (Heated electrospray ionization) probe. MS parameters were used as follows: spray voltage 3.5 kV and 3.2 kV for positive and negative modes, respectively; probe temperature 320 °C; sheath and auxiliary gases were 30 and 5 arbitrary units, respectively; full scan range: 70 m/z to 1050 m/z with settings of AGC target (3 × 10^6^) and mass resolution as Balanced and High (70,000). Data were recorded using Xcalibur 3.0.63 software (Thermo Scientific; Waltham MA, USA). Before analysis, Thermo Scientific Calmix solution was used as a standard to perform mass calibration in both ESI polarities and ubiquitous low-mass contaminants were used to apply lock-mass correction to each analytical run in order to enhance calibration stability. Parallel reaction monitoring (PRM) acquisition parameters: resolution 17,500, auto gain control target 2 × 10^5^, maximum isolation time 100 ms, isolation window m/z 0.4; collision energies were set individually in HCD (high-energy collisional dissociation) mode. Quality control samples were generated by taking equal volumes of each sample and pooling them, and subsequently analysing this mix throughout the run to assess the stability and performance of the system. Qualitative and quantitative analysis was performed using Xcalibur Qual Browser and Tracefinder 4.1 software (Thermo Scientific; Waltham MA, USA) according to the manufacturer’s workflows.

### Metabolite extraction and cell volume calculations

24 hours prior to the experiment, cells were seeded in 6 cm dishes in RPMI media containing 10 % dialysed FCS (3500 Da MWCO, PBS used for dialysis). An hour prior to treatment, the media were refreshed, and then changed again at t = 0 to RPMI with or without 11mM glucose; or to HBSS (H2969-500mL; Sigma Aldrich, St. Louis MS, USA) with or without supplemented amino acids as described in the text. Four technical replicate plates were used for each condition, and 2-3 plates for each cell line were used to count cells and measure mean cell diameter which was then used to determine cell volume in order to estimate intracellular concentrations. After 1 hour of treatment, plates were washed twice with ice-cold PBS, and 725 μl of dry-ice-cold methanol was used to quench the cells. Plates were scraped and contents were transferred to Eppendorf tubes on ice containing 180 μl H_2_O and 160 μl CHCl_3_. A further 725 μL methanol was used to scrape each plate and added to the same Eppendorf. Samples were vortexed and sonicated in a cold sonicating water bath 3 times for 8 mins each time. Extraction of metabolites was allowed to proceed at 4 °C overnight, before spinning down precipitated material and then drying down supernatant. To split polar and apolar phases, dried metabolites were resuspended in a 1:3:3 mix of CHCl_3_/ MeOH/ H_2_O (total volume of 350 μl). Polar metabolites in the aqueous phase were then analysed by LC-MS. To enable absolute quantification of metabolites of interest, known quantities of ^13^C-labelled versions of those metabolites were added into lysates, all purchased from Cambridge Isotopes (Tewksbury MA, USA). Previously determined cell numbers and volumes were then used to determine intracellular concentrations.

| Metabolite | Formula | Exact Mass | Pos. Mode m/z | Neg. Mode m/z | Mode used | RT (min) |
| --- | --- | --- | --- | --- | --- | --- |
| FBP | C<sub>6</sub>H<sub>14</sub>O<sub>12</sub>P<sub>2</sub> | 339.99611 | 341.00394 | 338.98829 | neg. | 13.42 |
| <sup>13</sup>C<sub>6</sub>-FBP | <sup>13</sup>C<sub>6</sub>H<sub>14</sub>O<sub>12</sub>P<sub>2</sub> | 346.01621 | 347.02404 | 345.00839 | neg. | 13.42 |
| PEP | C<sub>3</sub>H<sub>5</sub>O<sub>6</sub>P | 167.98241 | 168.99023 | 166.97458 | neg. | 13.58 |
| <sup>13</sup>C<sub>2</sub>-PEP | <sup>13</sup>C<sub>2</sub>C<sub>1</sub>H<sub>5</sub>O<sub>6</sub>P | 169.98911 | 170.99693 | 168.98128 | neg. | 13.58 |
| Ser | C<sub>3</sub>H<sub>7</sub>NO<sub>3</sub> | 105.0426 | 106.05043 | 104.03478 | pos. | 13.14 |
| <sup>13</sup>C<sub>3</sub>-Ser | <sup>13</sup>C<sub>3</sub>H<sub>7</sub>NO<sub>3</sub> | 108.05265 | 109.06048 | 107.04483 | pos. | 13.14 |
| Phe | C<sub>9</sub>H<sub>11</sub>NO<sub>2</sub> | 165.07899 | 166.08681 | 164.07116 | pos. | 9.7 |
| <sup>13</sup>C<sub>6</sub>-Phe | <sup>13</sup>C<sub>6</sub>H<sub>11</sub>NO<sub>2</sub> | 171.09909 | 172.10691 | 170.09126 | pos. | 9.7 |

### Targeted proteomics

#### Trypsin digestion

Cell extracts containing 50 μg of total protein were precipitated by adding six volumes of ice-cold acetone (pre-cooled to −20 °C). The samples were allowed to precipitate overnight at −20 °C and centrifuged at 8000 g, for 10 min at 4°C, to collect the pellet. The supernatant was carefully decanted and the residual acetone was evaporated at ambient temperature. The pellet was dissolved in 50 mM TEAB, reduced with 10 mM DTT and alkylated with 20 mM iodoacetamide. After alkylation, the proteins were digested with 1 μg of trypsin overnight at 37°C. After digestion, each sample was spiked with a mixture of five heavy labelled standards. For MS analysis, 1 μg of peptides were loaded onto 50-cm Easy Spray C18 column (Thermo Scientific; Waltham MA, USA).

#### Analysis of peptides by LC-tandem MS (LC-MS/MS)

Mass spectrometric analysis was performed using a Dionex U3000 system (SRD3400 degasser, WPS-3000TPL-RS autosampler, 3500RS nano pump) coupled to a QExactive electrospray ionisation hybrid quadruole-orbitrap mass spectrometer (Thermo Scientific; Waltham MA, USA). Reverse phase chromatography was performed with a binary buffer system at a flow rate of 250 nL/min. Mobile phase A was 5 % DMSO in 0.1 % formic acid and mobile phase B was 5 % DMSO, 80 % acetonitrile in 0.1 % formic acid. The digested samples were run on a linear gradient of solvent B (2 – 35 %) in 90 minutes, the total run time including column conditioning was 120 minutes. The nanoLC was coupled to a QExactive mass spectrometer using an EasySpray nano source (Thermo Scientific; Waltham MA, USA). The spray conditions were: spray voltage +2.1 kV, capillary temperature 250 °C and S-lens RF level of 55. For the PRM (parallel reaction monitoring) experiments, the QExactive was operated in data independent mode. A full scan MS1 was measured at 70,000 resolution (AGC target 3 × 10^6^, 50 ms maximum injection time, m/z 300-1800). This was followed by ten PRM scans triggered by an inclusion list (17,500 resolution, AGC target 2 × 10^5^, 55 msec maximum injection time). Ion activation/dissociation was performed using HCD at normalised collision energy of 28.

#### Data analysis of PRM

Peptides to be targeted in the PRM-MS analysis were selected previously by analysing trypsin digested cell extracts from the three cell lines of interest. Peptides providing a good signal and identification score representing the two PKM isoforms (PKM1/2) were selected for the analysis. The corresponding heavy isotope-labelled standards were synthesised in-house. The PRM method was developed for the QExactive using Skyline 4.1.0.18169. The heavy labelled peptide standards were used to create the precursor (inclusion) list. When measuring the abundance of PKM1/2 in the cell extracts signal extraction was performed on + 2 precursor ions for both heavy and light forms of the peptides. A peptide was considered identified if at least four overlapping transitions were detected. Quantitation was performed using MS2 XICs where the top three transitions were summed and used for quantitation. Data processing was performed with Skyline which was used to generate peak areas for both light and heavy peptides. Extracted ion chromatograms were visually inspected and peak boundaries were corrected and potential interferences removed. The data was subsequently exported in Excel to calculate absolute quantities of the “native” peptides and to determine reproducibility (CV %) of the measurements. See **Supplementary Table**. Data are available via ProteomeXchange with identifier PXD010334.

### Recombinant protein expression and purification

Allosteric hub mutant plasmids were generated through a single-step PCR reaction using hot-start KOD polymerase (Merck Millipore; Burlington MA, USA) and a pET28a-His-PKM2(WT) template plasmid (# 42515 AddGene; Cambridge MA, USA). Plasmids were sequence-verified by Sanger Sequencing (Source Bioscience; Nottingham, UK). 40 ng of pET28a-His-PKM2 (wild-type or mutant) was transformed into 50 μL *E. coli* BL21(DE3)pLysS (60413; Lucigen, Middleton WI, USA). Colonies were inoculated in LB medium at 37 °C and grown to an optical density of 0.8 AU (600 nm), at which point expression of the N-terminal His_6_-PKM2(WT) was induced with 0.5 mM isopropyl ß-D-1 thiolgalactopyranoside (Sigma Aldrich, St. Louis MS, USA). The culture was grown at 24 °C for 16-18 h. Cells were harvested by centrifugation and the pellet was re-suspended in cell lysis buffer (50 mM Tris-HCl pH 7.5, 10 mM MgCl_2_, 200 mM NaCl, 100 mM KCl and 10 mM imidazole) supplemented with the EDTA-free Complete protease inhibitor cocktail (Sigma Aldrich, St. Louis MS, USA). Cells were lysed by sonication at 4 °C. DNase was added at 1 μL/mL before centrifugation of the lysate at 20000 g for 1 hour at 4 °C. The supernatant (the water-soluble cell fraction) was loaded onto a HisTrap HP nickel-charged IMAC column (GE; Boston MA, USA) and was washed with five column-volumes of wash buffer [10 mM HEPES pH 7.5, 10 mM MgCl_2_, 100 mM KCl, 10 mM imidazole and 0.5 mM tris-2-carboxyethyl phosphine hydrochloride (TCEP; Sigma Aldrich, St. Louis MS, USA)]. After consecutive wash steps, the protein was eluted from the IMAC column with elution buffer buffer (10 mM HEPES pH 7.5, 10 mM MgCl_2_, 100 mM KCl, 250 mM imidazole and 0.5 mM TCEP). The N-terminal His_6_-epipope tag was cleaved with at 4 °C for ∼18 hours in cleavage buffer (50 mM Tris-HCl pH 8.0, 10 mM CaCl_2_) with recombinant bovine thrombin, immobilised on agarose beads. Purified recombinant PKM2 was eluted from the thrombin-agarose column. Affinity purification was followed by size-exclusion chromatography on a HiLoad 16/60 Superdex 200 pg column (28-9893-35; GE, Boston MA, USA) at 500 mL/min flow rate with protein storage buffer (10 mM HEPES pH 7.5, 10 mM MgCl_2_, 100 mM KCl, and 0.5 mM TCEP) at 4 °C. Eluted PKM2 was collected and concentrated to a final protein concentration of ∼7 mg/mL with centrifugal concentrating filters (Vivaspin 20, 10 kDa molecular-weight cut-off, 28-9323-60; GE, Boston MA, USA). Protein purity was assessed by SDS-PAGE. The final concentration of the protein was obtained by measuring the fluorescence absorbance spectrum between 240 nm and 450 nm. The concentration was calculated using a molar extinction coefficient of 29,910 M^−1^ cm^−1^ at 280 nm.

### Quantification of co-purified D-fructose-1,6-bisphosphate

Molar amounts of D-fructose-1,6-bisphosphate (FBP) that co-purified with recombinant PKM2 were measured using an aldolase enzyme assay that comprises three coupled enzymatic steps (**Supplementary Fig. 1a**). The reaction mixture contained 20 mM Tris-HCl pH 7.0, 50 μM NADH, 0.7 U/mL glycerol 3-phosphate dehydrogenase (G-3-PDH), 7 U/ml triose phosphate isomerase (TPI) and the supernatant of 5-50 μM purified recombinant PKM2 after heat-precipitation at 90 °C. G-3-PDH and TPI from rabbit muscle were purchased as a mixture (50017, Sigma Aldrich; St. Louis MS, USA). The reaction was initiated by adding between 0.008 – 0.016 U/ml rabbit muscle aldolase (A2714, Sigma Aldrich; St. Louis MS, USA) to a total reaction volume of 100 μL. Two molecules of NADH are oxidised for each molecule of FBP consumed. NADH oxidation was monitored over time at 25 °C in a 1 mL quartz cuvette (1 cm path-length) by measuring the NADH absorption signal at 340 nm using a Jasco V-550 UV-Vis spectrophotometer. For the assay calibration, known amounts of FBP from a powder stock were used instead of the heat-precipitated PKM2 supernatant.

### Measurement of PKM2 steady-state enzyme kinetics

Steady-state enzyme kinetic measurements of PKM2 were performed using a Tecan Infinite 200-Pro plate reader (Tecan, Männedorf Zürich, Switzerland). Initial velocities for the forward reaction (phosphoenolpyruvate and adenosine diphosphate conversion to pyruvate and adenosine triphosphate) were measured using a coupled reaction with rabbit muscle lactate dehydrogenase (Sigma Aldrich, St. Louis MS, USA). The reaction monitored the oxidation of NADH (*ε_340 nm_* = 6220 *M*^−1^*cm*^−1^) at 37 °C in a buffer containing 10 mM Tris-HCl pH 7.5, 100 mM KCl, 5 mM MgCl_2_ and 0.5 mM TCEP. Initial velocity versus substrate concentrations for phosphoenolpyruvate were measured in the absence and in the presence of allosteric ligands, in a reaction buffer containing 180 μM NADH and 8 U lactate dehydrogenase. Reactions were initiated by adding phosphenolpyruvate (PEP) at a desired concentration, with adenosine diphosphate (ADP) at a constant concentration of 5 mM. A total protein concentration of 5 nM PKM2 was used for all enzyme reactions, in a total reaction volume of 100 μL per well. Kinetic constants were determined by fitting initial velocity curves to Michaelis-Menten steady-state kinetic models.

### Measurements of FBP binding to PKM2

The affinity of PKM2 for FBP was measured by titrating small aliquots of a concentrated stock solution of FBP into 5 μM recombinant PKM2 and recording intrinsic fluorescence emission spectra of PKM2. Spectra were recorded using a Jasco FP-8500 spectrofluorometer with an excitation wavelength of 280 nm (bandwidth of 2 nm) and emission scanned from 290 nm to 450 nm (bandwidth of 5 nm) in a 0.3 cm path length quartz cuvette (Hellma Analytics; Muellheim, Germany) at 20 °C. The ratio of the emission intensities at 325 and 350 nm was plotted against the concentration of the titrant. Binding curves were fit to a model assuming a 1:1 binding stoichiometry (1 FBP molecule per monomer of PKM2) with a non-linear least squares regression fit of the following equation:

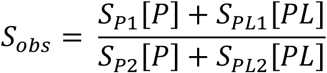

with [*P*] = [*P*_0_] – [*PL*] and [*PL*] calculated using

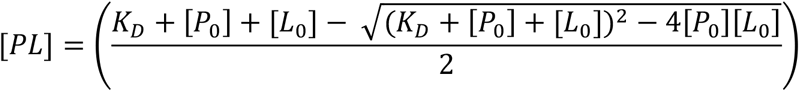

where the spectral signal *S_obs_* is the ratio of fluorescence emissions at wavelengths 1 (325 nm) and 2 (350 nm); *S_P1_, S_P2_, S_PL1_* and *S_PL2_* are the fluorescence extinction coefficients of the free protein *P* and the protein-ligand complex *PL* at wavelengths 1 and 2, respectively; [*P*_0_] and [*L*_0_] are total concentrations of protein and ligand, respectively; and *K_D_* is the apparent dissociation constant. The total free protein concentration ([*P*_0_]) was corrected by subtracting the percentage of protein pre-bound to co-purified FBP, as determined from the aldolase assay.

### Measurements of phenylalanine (Phe) and serine (Ser) binding to PKM2

The binding of Phe and Ser to PKM2 was measured using microscale thermophoresis (MST) on a Monolith NT.115 instrument (Nanotemper Technologies; Munich, Germany). First, PKM2 was fluorescently labelled with an Atto-647 fluorescein dye (NT-647-NHS; Nanotemper Technologies; Munich, Germany). 250 μL of 20 μM recombinant PKM2 was labelled with 250 μL of 60 μM dye in a buffer containing 100 mM bicarbonate pH 8.5 and 50% DMSO for 30 minutes at room temperature in the dark. Free dye was separated from labelled PKM2 using a NAP-5 20 ST size-exclusion column (GE; Boston MA, USA). Labelled PKM2, at a constant concentration of 30 nM, was titrated with either Phe (up to 5 mM) or Ser (up to 10 mM) in a buffer containing 10 mM HEPES pH 7.5, 100 mM KCl, 5 mM MgCl_2_, 0.5 mM TCEP and 0.1 % tween-20. Prior to each thermophoresis measurement, capillary scans were obtained to determine sample homogeneity. Binding curves were fit assuming a 1:1 stoichiometry.

### Analysis of the steady-state kinetics of PKM2 enzyme activity inhibition by Phe

In order to assign the mechanism with which Phe inhibition of PKM2 occurs, in the absence and in the presence of FBP, the dependence of the enzyme kinetic constants *K_M_*, *k_cat_* and 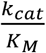 on the concentration of Phe were determined. Steady-state measurements of PKM2 enzyme activity (as described above) were performed by titrating the substrate PEP at several different concentrations of Phe and a constant concentration of 5 mM ADP. In order to investigate the allosteric K-type effect of Phe on enzyme affinity for its substrate PEP, a single-substrate-single-effector paradigm was assumed. Under equilibrium conditions the rate equation of the general modifier mechanism reveals apparent values of *K_M_* and *k_cat_*:

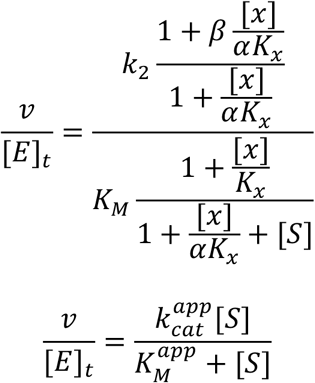

where [*E*]_*t*_ is the concentration of enzyme active sites, *x* is phenylalanine, *S* is the substrate (phosphoenolpyruvate), *K_x_* is the dissociation constant of the specific component of the enzyme mechanism, *α* is the reciprocal allosteric coupling constant and *β* is the factor by which the inhibitor affects the catalytic rate constant *k*_2_^51^. The kinetic constants 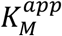, 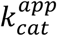 and 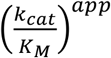 on the concentration of the modifier (X; phenylalanine) can then be written as follows^51^:

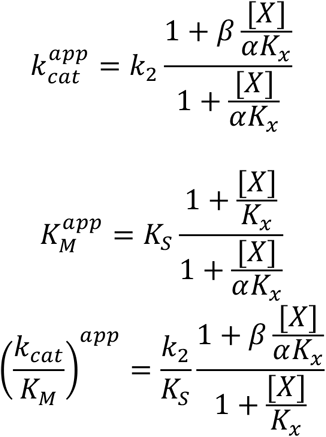

Rate curves measuring the dependence of [Phe] on the three kinetic constants were fit to the equations above to solve for the constants *K_S_, K_x_, α* and *β*. The kinetic mechanism was assigned based on the topology of rate-modifier mechanisms detailed by Baici^51^.

### Native Mass Spectrometry

PKM2 samples were buffer exchanged into 200 mM ammonium acetate (Fisher Scientific; Loughborough, UK) using Micro Bio-Spin 6 chromatography colums (Bio-Rad Laboratories; Hercules CA, US). Samples were diluted to a final protein concentration of between 5 μM and 20 μM depending on the experiment. Ligands were dissolved in 200 mM ammonium acetate and added to the protein prior to MS analysis.

Native mass spectrometry experiments were performed across three different instruments: a Ultima Global (Micromass; Wilmslow, UK) extended for high mass range, a modified Synapt G2 (Waters Corp, Wilmslow, UK) where the triwave assembly was replaced with a linear drift tube, and a Synapt G2-Si. Samples were analysed in positive ionization mode using nano-electrospray ionization, in which the sample is placed inside a borosilicate glass capillary (World Precision Instruments; Stevenage, UK) pulled in-house on a Flaming/Brown P-1000 micropipette puller (Sutter Instrument Company; Novato CA, USA) and a platinum wire is inserted into the solution to allow the application of a positive voltage. All voltages used were kept as low as possible to achieve spray while keeping the protein in a native-like state. Typical conditions used were capillary voltage of ∼1.2 kV, cone voltage of ∼10 V and a source temperature of 40 °C.

### Ion mobility mass spectrometry (IM-MS)

IM-MS measurements were performed on an in-house modified Synapt G2 in which the original triwave assembly was replaced with a linear drift tube with a length of 25.05 cm. Drift times were measured in helium at a temperature of 298.15 K and pressure of 1.99-2.00 torr. Conditions were kept constant across each run. Measurements were performed at least twice for each sample and averaged. Mobilities for all charge states were converted into rotationally averaged collision cross sections (^DT^CCS_He_) using the Mason-Schamp equation^78^, and further converted into a single global collision cross section distribution per species which all charge states contribute towards in proportion to their intensity in the mass spectrum.^79^.

### Surface-Induced Dissociation (SID)

SID was performed on a prototype instrument at Waters Corp (Wilmslow, UK). Briefly, sample was ionized via nano-ESI and the species of interest was mass selected in the quadrupole. Ions were then accelerated towards the gold surface of an SID device and underwent a single, high energy collision. Fragments were refocused and mass analysed.

### Molecular dynamics simulations in explicit solvent

Molecular dynamics (MD) simulations of tetrameric human PKM2 were performed with the GROMACS 5.2 engine^80^, using the Gromos 53a6 force field parameter sets^81^ and SPC-E water molecules. The input coordinates for PKM2^apo^ were extracted from the Protein Data Bank crystal structure 3bjt^46^ and coordinates for PKM2^FBP^ were extracted from the crystal structure 3u2z^20^. The force-field parameters for FBP were determined using a quantum mechanical assignment of the partial charges using the ATB server^82^. Structures were prepared as previously described^72^. Briefly, structures were solvated in a dodecahedral period box, such that the distance between any protein atom and the periodic boundary was a minimum of 1.0 nm. The system charge was neutralised by adding counter ions to the solvent (Na^+^ and Cl^−^). Equations of motion were integrated using the leap-frog algorithm^83^ with a 2 fs time step. The system was equilibrated for 5 ns in the NVT ensemble at 300 K and 1 bar. This was followed by a further 5 ns equilibration in the NPT ensemble. Following equilibration, five replicate production run simulations were performed for 400 ns under constant pressure and temperature conditions, 1 bar and 300 K. Temperature was regulated using the velocity-rescaling algorithm^84^, with a coupling constant of 0.1. Covalent bonds and water molecules were restrained with the LINCS and SETTLE methods, respectively. Electrostatics were calculated with the particle mesh Ewald method, with a 1.4 nm cut-off, a 0.12 nm FFT grid spacing and a four-order interpolation polynomial for the reciprocal space sums.

### Prediction of allosteric hub residues with AlloHubMat

MD trajectories were coarse-grained with the M32K25 structural alphabet using the GSAtools^63^. From the fragment-encoded trajectory, the mutual information between each combination of fragment positions 〈*I^n^*(*C_i_; C_j_*)〉 was determined for multiple replicate 400 ns trajectories for PKM2^apo^ and PKM2^FBP^. Each trajectory was sub-divided into 20 nonoverlapping blocks with an equal time length of 20 ns each. For each trajectory block (*B*), correlated conformational motions for all pairs of fragments (*i,j*) were calculated as the normalised mutual information between each fragment pair in the fragment-encoded alignment 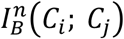:

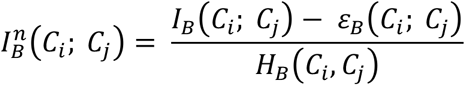

where the columns of the structural fragment alignment are given by *C_i_* and *C_j_, I_B_*(*C_i_; C_j_*) is the mutual information, *ε_B_*(*C_i_; C_j_*) is the expected finite size error and *H_B_*(*C_i_,C_j_*) is the joint entropy^54^. In this framework, the mutual information is given by:

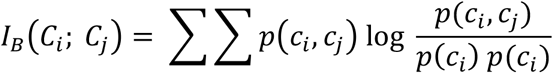

where the two columns in the structural alphabet alignment *C_i_* and *C_j_* are random variables with a joint probability mass function *p*(*c_i_, c_j_*), and marginal probability mass functions *p*(*c_i_*) and *p*(*c_j_*). The joint entropy *H*(*C_i_; C_j_*) is defined as:

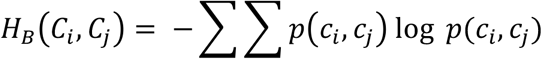

The discrete mutual information calculated for finite state probabilities can be significantly affected by random and systematic errors. In order to account for this, we subtract an error term *ε_B_*(*C_i_; C_j_*) given by:

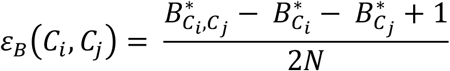

where *N* is the sample size and 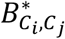, 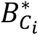 and 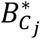 are the number of states with non-zero probabilities for *C_i_C_j_, C_i_* and *C_j_*, respectively^85^.

With the goal of identifying conformational sub-states and their respective probabilities from each of the trajectories, eigenvalue decomposition was used to compute the geometric evolution of the protein backbone correlations (given by the mutual information between structural-alphabet fragments). The elements of the mutual information matrix are proportional to the square of the displacement, so the square root of the matrix is required to examine the extent of the matrix overlap:

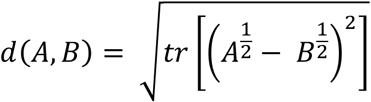

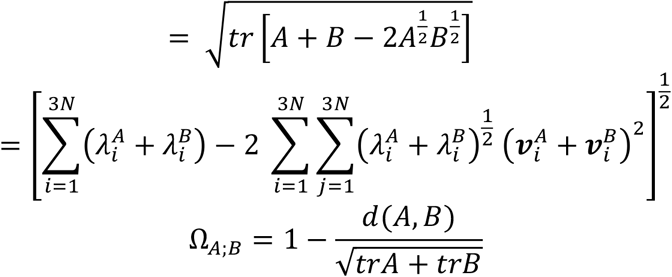

where 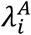 and 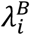 denote the eigenvalues and 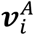 and 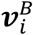 the eigenvectors of mutual information matrices *A* and *B, N* is the number of structural alphabet fragments. The covariance matrix overlap (Ω_*A;B*_) ranges between 0 when matrices *A* and *B* are orthogonal, and 1 when they are identical.

Using the approach detailed above, the covariance overlap (Ω_*A;B*_) was determined from time-contiguous mutual information matrices extracted from non-overlapping trajectory blocks. Conformational sub-states were identified as containing a high degree of similarity between time-contiguous mutual information matrices. Using this approach, an ensemble-averaged mutual information matrix for PKM2^apo^ and for PKM2^FBP^ was determined by averaging over all mutual information matrices identified from each conformational sub-state of the multiple replicate simulations of both liganded states.

The above approach made it possible to subtract the mutual information matrices of correlated motions of the holo-from the apo-state. A difference mutual information matrix was constructed by subtracting the ensemble-averaged matrix of PKM2^FBP^ from PKM2^apo^. Allosteric hub fragments (AlloHubFs) were identified from this difference mutual information matrix, as those fragments with the highest log_2_-fold change in the coupling strength and a p-value associated with the change of less than 0.01.

### Estimation of the configurational entropy from explicit solvent MD simulations

The configurational entropy of MD trajectories of PKM2^apo^ and PKM2^FBP^ were estimated using a formulism proposed by Schlitter^86^. For a classical-mechanical system the configurational entropy is given by:

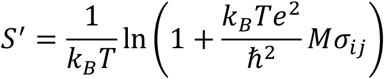

where

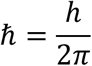

k_B_ is the Boltzmann constant, T is temperature, e is Euler’s number, *h* is Plank’s constant and *Mσ_ij_* is the mass-weighted covariance matrix of the form:

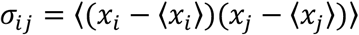

where x_i_ and x_j_ are positional coordinates of the MD trajectory in Cartesian space.

### Theoretical collision cross section calculations

Structural models of the 15+ monomer, 23+ A-A’ and C-C’ dimers and 33+ tetramers were generated from the PDB crystal structure 3bjt^46^ and simulated *in vacuo* using the OPLS-AA/L force-field parameter set^87^. Systems were minimised using the Steepest Descent algorithm for 5 × 10^6^ steps, with a step size of 1 J mol^−1^ nm^−1^ and a maximal force tolerance of 100 kJ mol^−1^ nm^−1^. Next, systems were equilibrated at consecutively increasing temperatures (100 K, 200 K and 300 K) each for 5 ns, with the Berendsen temperature coupling method^83^ and an integration step size of 1 fs. Following successful equilibration, 10 ns simulations were performed with an integration step size of 2 fs. Pressure coupling and electrostatics were turned off. Temperature was held constant at 300 K using the Berendsen coupling method.

The most prevalent structures were extracted using the GROMOS clustering algorithm^68^. Theoretical collision cross sections were calculated for each clustered structure, using the projection approximation method (as outlined in Ruotolo, B.T. *et al*. ^88^) and using the exact hard sphere scattering model (as implemented in the EHSSrot software^89^).

### Circular dichroism (CD) spectroscopy

Far-UV circular dichroism (CD) spectra were recorded using a JASCO J-815 spectrometer (Jasco; Oklahoma City, OK USA) from 200 nm to 260 nm with 300 μL of 0.2 mg/mL PKM2 ion a quartz cuvette with a path length of 0.1 cm. Measurements were performed at a constant temperature of 20 °C. Raw data in units of mdeg were converted to the mean residue CD extinction co-efficient, in units of M^−1^ cm^−1^:

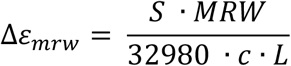

where *c* is the protein molar concentration, *L* is the path length (cm), *S* is the raw measurement of CD intensity (in units of mdeg) and *MRW* is the molecular weight of the protein divided by the number of amino acids in the protein.

### Calculation of the allosteric coupling co-efficient for wild-type PKM2 at the AlloHub mutants

Enzyme activity measurements of the PKM2(WT) and the AlloHubMs were performed at 37 °C using a lactate dehydrogenase assay, as previously described. Initial velocities were measured over a range of phosphoenolpyruvate concentrations, with a constant concentration of 5 mM ADP. Measurements were repeated following pre-incubation of the PKM2 variant with saturating concentrations of FBP (2 μM for all variants, with the exception of R489L which was incubated with 50 mM FBP). The allosteric coupling constant (*Q*) was calculated to determine the coupling between FBP binding and catalysis, as previously described^50,58,90^:

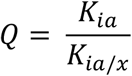

where *K_ia_* and *K_ia/x_* are equilibrium dissociation constants for the binding of the substrate (*a*) in the absence or presence, respectively, of the allosteric effector *x*. When *Q* > 1, there is positive allosteric coupling between the binding of *x* to the protein and the binding of A to the substrate binding pocket. Conversely, when *Q* < 1, there is negative coupling between the A and x sites. Measurements were repeated after addition of 400 μM Phe to the protein variants that had been pre-incubated with FBP, and activity was measured over a range of substrate concentrations. This facilitated the calculation of the coupling constants between the Phe, FBP and substrate binding sites.

### Code Availability

The computer code used in this study is available from the corresponding authors upon reasonable request.

### Data Availability

The data that support the findings of this study are available from the corresponding authors upon reasonable request. Accession codes and relevant web links can be found in the respective legends and methods sections.

